# Distinct evolutionary trajectories of neuronal and hair cell nicotinic acetylcholine receptors

**DOI:** 10.1101/621342

**Authors:** Irina Marcovich, Marcelo J. Moglie, Agustín E. Carpaneto Freixas, Anabella P. Trigila, Lucia F. Franchini, Paola V. Plazas, Marcela Lipovsek, Ana Belén Elgoyhen

## Abstract

The expansion and pruning of ion channel families has played a crucial role in the evolution of nervous systems. Remarkably, with a highly conserved vertebrate complement, nicotinic acetylcholine receptors (nAChRs) are unique among ligand-gated ion channels in that members of the family have distinct roles in synaptic transmission in non-overlapping domains, either in the nervous system, the inner ear hair cells or the neuromuscular junction. Here, we performed a comprehensive analysis of vertebrate nAChRs sequences, single cell expression patterns and comparative functional properties of receptors from three representative tetrapod species. We show that hair cell nAChRs underwent a distinct evolutionary trajectory to that of neuronal receptors. These were most likely shaped by different co-expression patterns and co-assembly rules of component subunits. Thus, neuronal nAChRs showed high degree of coding sequence conservation, coupled to greater co-expression variance and conservation of functional properties across tetrapod clades. In contrast, hair cell α9α10 nAChRs exhibited greater sequence divergence, narrow co-expression pattern and great variability of functional properties across species. These results point to differential substrates for random change within the family of gene paralogs that relate to the segregated roles of nAChRs in synaptic transmission.

**Significance statement:** Our work exploits several peculiarities of the family of vertebrate nicotinic acetylcholine receptors (nAChRs) to explore the evolutionary trajectories of a ligand-gated ion channel family. By performing a comprehensive comparative analysis of nAChR subunits coding sequences, single cell expression patterns and functional properties we found a contrasting evolutionary history between nAChRs with widespread expression in the nervous system compared to those with isolated expression in the inner ear. Evolutionary changes were focused on differences in co-expression and co-assembly patterns for the former and coding sequences in the latter. This multidisciplinary approach provides further insight into the evolutionary processes that shaped the nervous and sensory systems of extant animals.

## INTRODUCTION

The superfamily of pentameric ligand-gated ion channels (LGIC) has an extended evolutionary history and is present in all three life domains (1–3). Extant members of the superfamily include eukaryote Cys-loop receptors that respond to acetylcholine (ACh), γ-aminobutyric acid (GABA), glycine or serotonin, invertebrate LGICs that respond to GABA, glutamate, ACh, serotonin and histamine, as well as prokaryote pH and GABA sensitive receptors, among others (1, 3–8). Nicotinic acetylcholine receptors (nAChRs) are a major branch of the Cys-loop LGIC superfamily (7, 9). To date, 17 different nAChR subunits have been described in the main vertebrate clades: α1-α10, β1-β4, δ, ε and γ (7, 8). These paralogous genes are proposed to derive from 5 paralogons, with the entire complement of extant subunits present in the vertebrate ancestor (10). Vertebrate nAChRs are non-selective cation channels that participate in numerous processes, most prominently neuromuscular junction (11, 12), inner ear efferent (13) and neuronal (14, 15) synaptic transmission.

Functional nAChRs are homomeric, comprising five identical subunits, or heteromeric, formed by at least two different subunits (7, 15, 16). The rules that govern the combinatorial assembly of functional nAChRs are complex and for the most part unknown. Some nAChR subunits can combine with other numerous, albeit specific, subunits. In contrast, some subunits can only form functional receptors with a limited subset (7, 16). This segregation has given rise to subgroups of vertebrate nAChRs named for their initially described tissue of origin and main functional location. Thus, neuronal nAChRs are formed by multiple combinations of α2-α7 (and α8 in non-mammals) and β2-β4 subunits, comprising a wide combinatorial range with alternative stoichiometries (15–17). Muscle receptors show tighter co-assembly rules. They have a typical α1_2_β1γδ (or α1_2_β1εδ) stoichiometry and do not co-assemble with non-muscle subunits (16, 18). Finally, the hair cell nAChR has a very strict co-assembly constraint, being formed exclusively by α9 and α10 subunits (19–21). While α9 subunits can form functional homomeric receptors (22, 23), these are unlikely to play a major role in inner ear hair cells in vivo (24). Although the α9 and α10 subunits were initially classified as a neuronal, the α9α10 receptor does not appear to be functionally present in the brain (25). A consequence of the differences in co-assembly rules between the three subgroups of nAChRs is that, while muscle cells can toggle between at least two receptor variants and neurons are capable of expressing a great diversity of nAChRs, hair cells express only one type.

The complement of nAChR subunits is highly conserved across vertebrates and even more so in tetrapods (1, 10). This suggests a high gene family-wide negative selection pressure for the loss of paralogs and highlights the functional relevance of each individual subunit across the vertebrate clade. However, given the major differences in co-expression patterns and co-assembly rules that distinguish the subgroups of nAChRs, in particular the contrast between neuronal and hair cell receptors, distinct evolutionary trajectories most likely took place in different members of the family. On the one hand, if only one (functionally isolated) receptor is expressed by a given cell type, then selection pressure could have acted on stochastic changes of the coding sequence of component subunits that altered receptor function. Such changes would not have resulted in deleterious effects on other nAChRs given the restricted expression pattern of the subunits. The aforesaid process could have dominated the evolutionary history of the hair cell α9α10 receptor. On the other hand, widely expressed subunits that co-assemble into functional receptors with multiple others have been most probably subjected to strong negative selection pressure. Changes in the coding sequence that lead to changes in functional properties may have had deleterious effects on alternative receptor combinations expressed in different cell types. In this case, functional diversification could have arisen from stochastic changes in the expression patterns of receptor subunits, resulting in a given cell changing the subtype of receptor it expresses, while preserving individual subunit functionality. Such processes could have dominated the evolutionary history of neuronal nAChRs. These contrasting theoretical scenarios bring forward three predictions for the evolutionary history of nAChRs across the vertebrate clade. First, isolated (hair cell) subunits may show coding sequence divergence while widespread (neuronal) subunits a high degree of coding sequence conservation. Second, isolated subunits may show low co-expression variation while widespread subunits a great variability in co-expression patterns. Finally, isolated receptors may present divergent functional properties across species resulting from changes in coding sequences, while widespread receptors may show highly conserved functional properties when formed by the same subunits.

To test these predictions, we studied the molecular evolution and the variability in expression pattern, coupled to co-assembly potential, of vertebrate nAChR subunits. Furthermore, we performed a comprehensive comparative analysis of the functional properties of the hair cell (α9α10) and the two main neuronal (α7 and α4β2) nAChRs from three representative species of tetrapods. In close agreement with the aforementioned predictions, neuronal subunits showed high degree of sequence conservation and great variability in co-expression patterns. Additionally, the functional properties of both α7 and α4β2 receptors did not differ across tetrapods. In contrast, the isolated hair cell α9α10 nAChRs showed the highest degree of sequence divergence and no co-expression variability. Moreover, the functional properties of α9α10 receptors from representative species of tetrapods showed unprecedented divergence, potentially echoing the evolutionary history of vertebrate auditory systems. In summary, here we present strong evidence supporting the notion that, within the family of paralog genes coding for nAChR subunits, receptors belonging to different subgroups (hair cell or neuronal) underwent different evolutionary trajectories along the tetrapod lineage that were potentially shaped by their different co-expression patterns and co-assembly rules. Furthermore, we propose that it is the difference in the most probable substrate for random change and subsequent selection pressure that separate the contrasting evolutionary histories of nAChRs subgroups.

## RESULTS

### Amino acid sequence divergence and co-expression patterns differentiate neuronal from hair cell nAChR subunits

The comparative analysis of gene paralogs provides the opportunity to test predictions about the evolutionary history of a gene family. To study the degree of conservation of coding sequences in subgroups of nAChRs we performed an exhaustive evaluation of sequence divergence of vertebrate nAChR subunits. The analysis included amino acid sequences from 11 species of birds, reptiles and amphibians, all groups that were notably under-represented in previous work (1, 26–30) and are of particular importance to help resolve differences between clades. Overall, we analysed 392 sequences from 17 nAChR subunits belonging to 29 vertebrate species (Table S1). Based on sequence identity, the family of nAChR subunits can be split into four groups: α, non-α, α7-like and α9-like (Fig. 1 and S1-S2). As previously reported with smaller datasets (26, 28), the present extended analysis showed that α10 subunits are unique in presenting a segregated grouping of orthologues with non-mammalian α10 subunits as a sister group to all α9 subunits, and mammalian α10 subunits an outgroup to the α9/non-mammalian α10 branch (Fig. 1). This may relate to the overall low %seqID of all vertebrate α10 subunits, coupled to high sequence conservation within individual clades (Table S2). Interestingly, a close inspection of the cumulative distribution of within and between-clade %seqID suggests that both α9 and α10 hair cell subunits may have accumulated a high proportion of their overall aminoacid changes at the mammalian/sauropsid split, while sequence divergence in neuronal subunits appears distributed throughout the phylogeny (Fig S3). To further test this, we searched for site-specific evolutionary shifts in aminoacid biochemical state between clades ((31) and SI methods) and found that both α9 and α10 subunits have a high number of sites showing potential functionally significant aminoacid changes when comparing the mammalian vs sauropsid clades (Fig. 1B – red bars and Table S3). In contrast, neuronal subunits fail to show between-clade functional divergence at the sequence level (Fig. 1B and Table S3). Overall, the extended molecular evolution analysis of nAChR subunits showed that while neuronal subunits were highly conserved across all vertebrate clades, α9 and α10 hair cell subunits showed a greater degree of between-clade sequence divergence that differentiates mammalian and sauropsid subunits.

**Figure 1.**
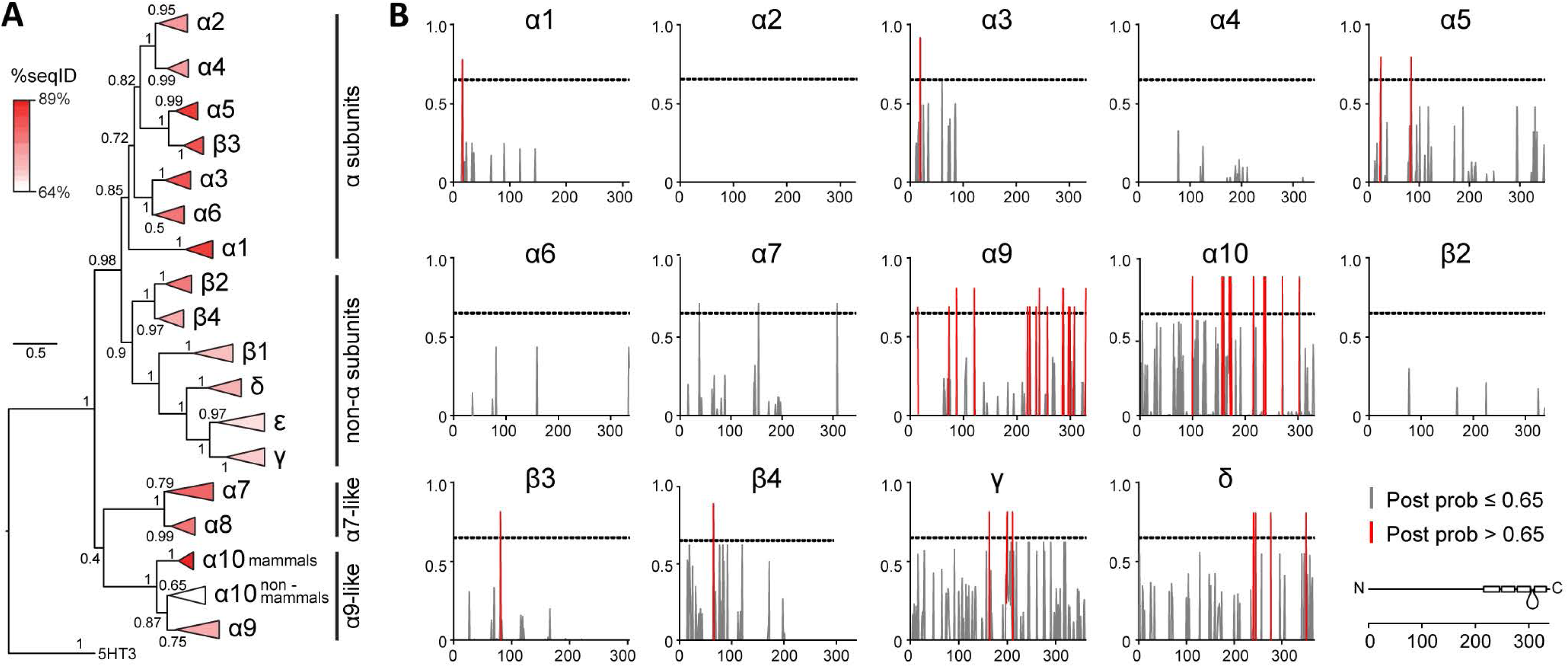
Hair cell nAChR subunits show greater sequence divergence than neuronal subunits. **A.** Phylogenetic relationships between vertebrate nicotinic subunits. The branches corresponding to the same subunits of different species were collapsed up to the node at which one subunit separates from its closest neighbour. The complete tree is shown in Fig. S1. Triangles length denotes the divergence on sequence identity from the subunit node. Triangles were coloured according to the average percentage of sequence identity between all pairs of sequences (%seqID, Table S2) within the branch. Numbers in branches indicate the bootstrap value obtained after 1,000 replicates. Scale bar indicates the number of amino acid substitutions per site. B. Posterior probabilities for type II functional divergence between mammalian and sauropsid clades, for each site along individual nAChR subunits. Grey lines, posterior probability ≤ 0.65. Red lines, posterior probability < 0.65. Bottom right, diagram of a nAChR subunit extracellular, 4 transmembrane and intracellular domains along aminoacid position.

The capability to co-assemble into functional receptors and the co-expression of nAChR subunits within a given cell delineate the complement of receptors that shape its nicotinic ACh response. Numerous heterologous expression experiments and subunit-specific pharmacological studies have outlined a comprehensive repertoire of functionally validated pentameric assemblies (Table S6 and references therein). However, no systematic gene expression analysis that explores the potential spectrum of nAChRs in a given cell type has been performed. In order to evaluate co-expression patterns, we performed a meta-analysis of gene expression data from 10 recent single-cell transcriptomic studies (Table S4). First, from the gene expression matrices we identified the repertoire of nAChR subunits detected in each single cell (Fig. S4). Subsequently, we used a Bayesian approach (32) to estimate the likelihood of a gene being expressed at any given average level in a given cell type (Table S5, Fig. 2 and Fig. S5). We next combined this data with the catalogue of validated nAChR pentamers (Table S6) and outlined the potential complement of pentameric receptors present in each cell type, by identifying the subunit combinations that are present within a 10-fold, 100-fold or 1000-fold range of expression level or altogether absent (Fig. S6A). As expected, neurons express a range of neuronal nAChR variants with major neuronal types identified by their well characterised nAChRs. For example, visceral motor neurons from thoracic sympathetic ganglia express receptors containing α3 and β4 subunits, while cortical neurons express receptors containing α4, β2 and α7 subunits. Ventral midbrain dopaminergic neurons express high levels of the α6 subunit, together with β2 and β3 subunits and variable levels of α3, α4 and α5 subunits. Receptors containing the α2 subunit are observed in different types of retinal neurons (Fig. 2). Also, GABAergic, glutamatergic and dopaminergic neurons from hypothalamus, ventral midbrain and/or sympathetic ganglia present noticeably different potential complements of nAChRs (Fig. S6A). Differences in potential nAChRs composition can be observed even between closely related cell types. For example, two subtypes of cortical pyramidal neurons that differ on their projection targets (33) show a significant difference in the expression level of the α5 subunit (Fig. S6B), indicating they could differ on the ratio of α4β2/α4α5β2 receptors on the plasma membrane. Finally, inner ear hair cells co-express high levels of α9 and α10 subunits (Fig. 2 and S6C). Taken together the analysis of aminoacid sequences and co-expression patterns indicates that, whereas the α9 and α10 subunits have a restricted expression pattern, they also have the highest clade-specific sequence divergence. On the contrary, neuronal nAChRs present higher sequence conservation together with a widespread expression pattern.

**Figure 2.**
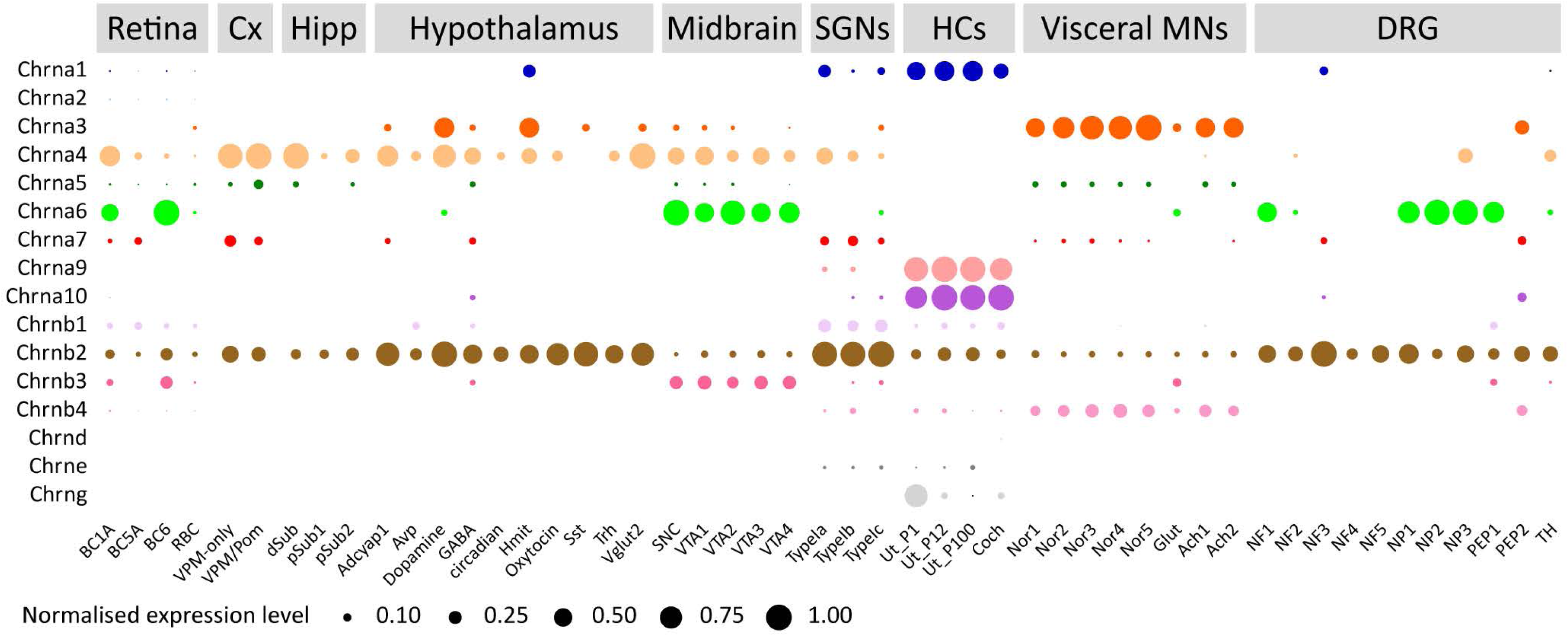
Hair cell nAChR subunits are co-expressed in inner ear hair cells, while neuronal subunits show widespread and variable co-expression patterns. Normalised mean expression level for nAChR subunits across mouse neuronal and sensory cell types. Circle sizes indicate the mean expression level for each cell type, normalised to the highest value observed within each dataset. For detailed explanations of individual cell types refer to main text, SI methods section or the original publications.

### Divergence of biophysical properties differentiates neuronal from hair cell nAChRs

The restricted co-expression pattern of hair cell α9 and α10 nAChR subunits (Fig. 2) and their exclusive co-assembly with each other (19–21), together with their higher level of between-clade sequence divergence (Fig. 1 and S3), indicate the potential for functional innovations across clades through coding sequence changes. In contrast, the high conservation of neuronal subunits (Fig. 1 and Table S3) may have resulted in highly conserved functional properties of neuronal receptors assembled from the same subunits. We experimentally tested this prediction by performing a comprehensive side-by-side comparison of the functional properties of the two main neuronal, α4β2 and α7 nAChRs, and the hair cell α9α10 nAChR from three representative species of tetrapod clades. To this end, we injected *Xenopus laevis* oocytes with the corresponding cRNAs and characterised the biophysical properties of ACh responses.

Oocytes injected with rat (*Rattus norvegicus*), chicken (*Gallus gallus*) or frog (*Xenopus tropicalis*) α4 and β2 cRNAs responded to ACh and showed the characteristic two-component ACh dose-response curves that correspond to the well described (α4)_3_(β2)_2_ and (α4)_2_(β2)_3_ prevalent stoichiometries (Fig. 3A, Table S8 and (34)). Oocytes injected with rat, chicken or frog α7 cRNA responded to ACh with similar apparent affinities (Fig. 3A, Table S8). Finally, rat, chicken and frog α9 and α10 subunits formed functional heteromeric α9α10 nAChRs that responded to ACh in a concentration-dependent manner (Fig. 3A and (19, 28)). The frog α9α10 receptor exhibited a significantly higher apparent affinity than its amniote counterparts (p = 0.0026 (vs rat), p = 0.0060 (vs chick) – Fig. 3A and Table S8).

**Figure 3.**
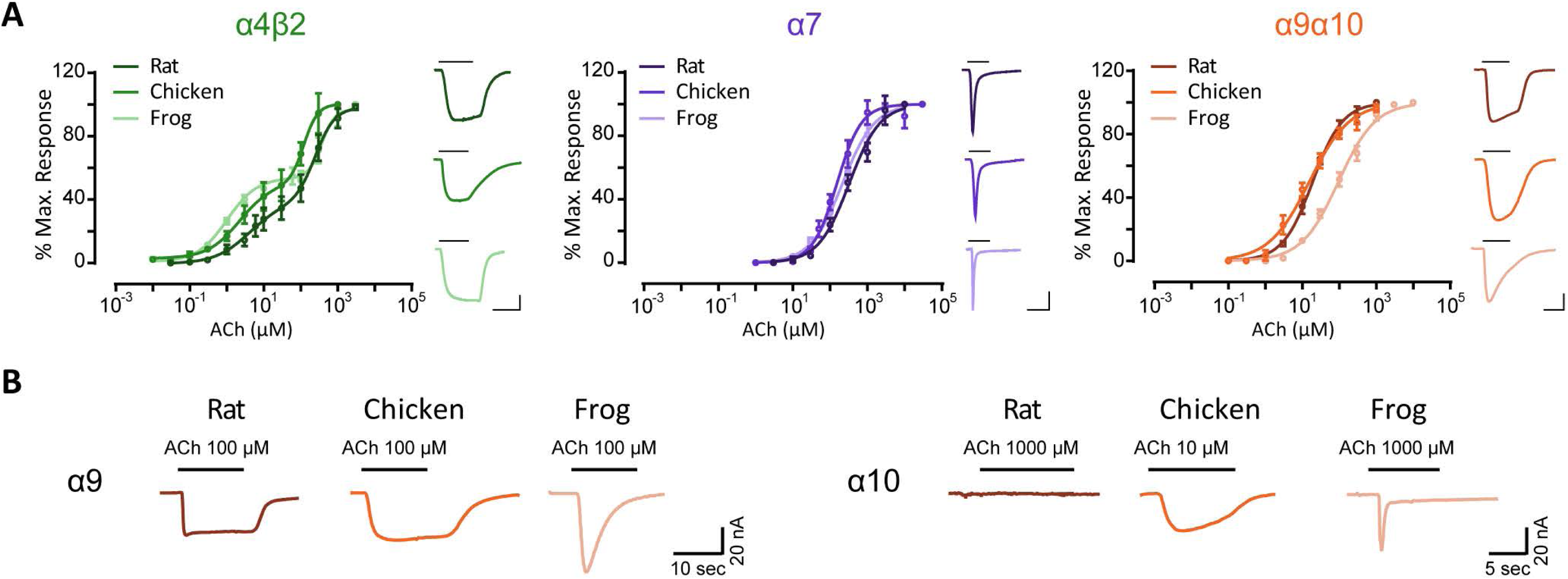
Hair cell nAChRs show differences in ACh apparent affinity, while neuronal nAChRs have similar ACh sensitivity. **A.** Concentration-response curves for neuronal α4β2 and α7 nAChRs and hair cell α9α10 nAChRs from three tetrapod species. Values are mean ± S.E.M. Lines are best fit to the Hill equation (n = 4-9). Representative responses evoked by 10 μM (α4β2, rat and chick α9α10) or 100 μM (α7, frog α9α10) ACh are shown next to their respective plots. Scale bars: α4β2: 100 nA, 10 sec; α7: 50 nA, 5 sec; α9α10: 50 nA, 10 sec. **B.** Representative responses evoked by ACh in oocytes injected with rat, chicken or frog homomeric α9 and α10 subunits (n=2–20).

While the α4 and β2 subunits participate exclusively in heteropentameric receptor assemblies, rat α9 (Fig. 3B and (22)), chicken α9 (Fig. 3B and (23))and frog α9 (Fig. 3B) subunits assembled into functional homomeric receptors. In contrast, rat α10 subunits cannot form functional receptors on their own (Fig. 3B and (19)). Surprisingly, both chicken α10 (Fig. 3B and (23)) and frog α10 (Fig. 3B) subunits assembled into functional homomeric receptors.

A defining feature of nAChRs is their desensitisation upon prolonged exposure to the agonist (35). Rat and chicken neuronal α4β2 receptors are characterised by slow desensitisation kinetics (28, 36). As shown in Fig. 4, 70-80% of the maximum current amplitude remained 20 seconds after the peak response to 100 μM ACh. Similarly, frog α4β2 receptors depicted slow desensitisation profiles, with no significant differences when compared to that of rat (p = 0.7326) and chicken α4β2 (p = 0.7204) nAChRs (Table S8 and Fig. 4). The frog α7 nAChRs showed fast desensitisation with 2-3% of current remaining 5 seconds after the peak response to 1 mM ACh, similar to that of rat α7 (p = 0.3743) and chicken α7 (p = 0.2496) nAChRs (Table S8 and Fig. 4).

**Figure 4.**
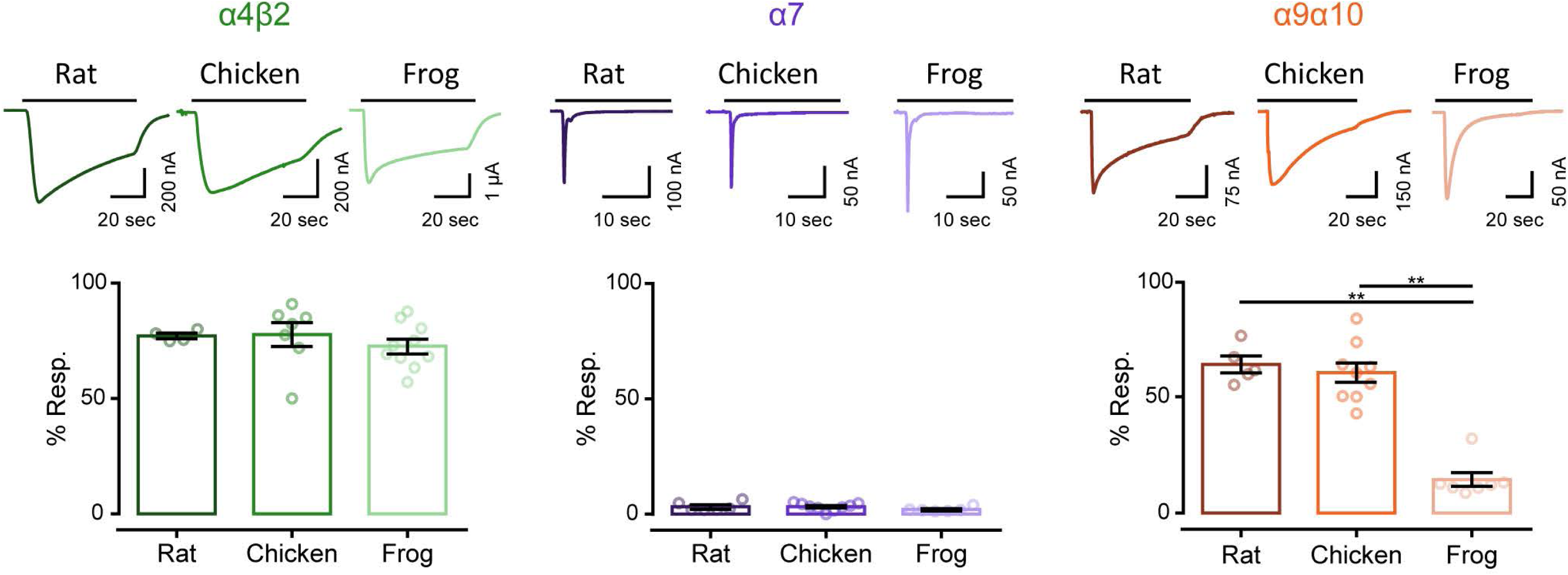
Hair cell nAChRs differ in their desensitization patterns, while neuronal receptors show similar profiles. *Top panels*. Representative responses of α4β2, α7 and α9α10 nAChRs to a 60 seconds (for α4β2 and α9α10) or 30 seconds (for α7) application of 100 μM ACh for all α4β2 and amniotes α9α10, and 1 mM ACh for all α7 and frog α9α10 nAChRs. *Bottom panels*. Percentage of current remaining 20 seconds (α9α10 and α4β2) or 5 seconds (α7) after the peak response, relative to the maximum current amplitude elicited by ACh. Bars represent mean ± S.E.M., open circles represent individual oocytes (n = 4-10). \*\**p*<0.01, One-Way ANOVA followed by Dunn’s test (α4β2 nAChRs) or Kruskal-Wallis followed by Holm Sidak’s test (α7 and α9α10 nAChRs).

The conserved desensitisation profiles observed for both types of neuronal receptors was in stark contrast with that of α9α10 receptors. While rat and chicken α9α10 receptors showed similar and somewhat slow desensitization, with 60-65% of current remaining 20 seconds after the peak response to ACh (p = 0.9999), frog α9α10 nAChRs exhibited significantly higher desensitization when compared to rat α9α10 (p=0.0051) and chicken α9α10 (p=0.0042) receptors (Fig. 4 and Table S7). Neuronal nAChRs are potentiated by extracellular divalent cations such as Ca^2+^ via an allosteric and voltage-independent mechanism (37, 38). The rat α9α10 nAChR, on the other hand, is both potentiated and blocked by physiological concentrations of extracellular divalent cations (39). Blockage occurs in the milimolar range, is voltage dependent and proposed to occur as a result of calcium permeation (39). To perform a comparative analysis of calcium modulation on rat, chicken and frog α4β2, α7 and α9α10 receptors, responses to near-EC_50_ concentrations of ACh were recorded in normal Ringer’s solution at a range of Ca^2+^ concentrations and normalised to the response at 1.8 mM Ca^2+^. For the neuronal α4β2 and α7 receptors from all three species, a similar potentiation pattern was observed, with increasingly higher responses to ACh at greater Ca^2+^ concentrations (Fig. 5 – top and middle panels). In contrast, responses of α9α10 nAChRs from rat, chicken and frog exhibited differential modulation by extracellular Ca^2+^. As previously reported for the rat α9α10 receptor, responses to ACh were potentiated and blocked by extracellular Ca^2+^ (Fig. 5 - bottom left panel and (39)). The chicken α9α10 receptor also showed peak potentiation of the ACh response at 0.5 mM extracellular Ca^2+^. However, no evident block was observed at higher concentrations of the cation (Fig. 5 - bottom middle panel). Finally, the frog α9α10 receptor showed potentiation of ACh responses at all Ca^2+^ concentrations assayed, with maximal responses detected at 3 mM Ca^2+^ (Fig. 5 - bottom right panel).

**Figure 5.**
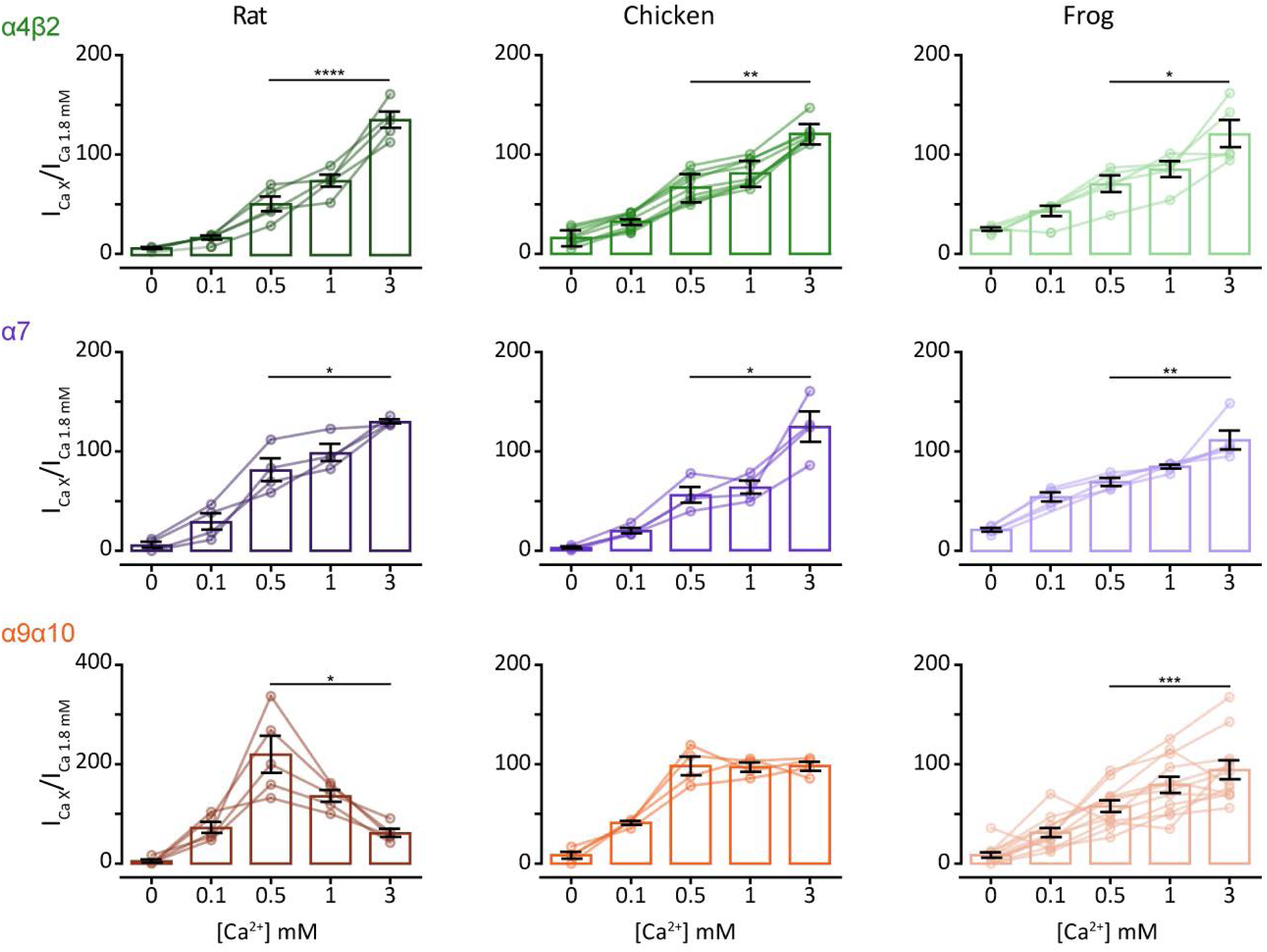
Extracellular Ca^2+^ potentiates neuronal nAChRs but differentially modulates α9α10 nAChRs. ACh response amplitude as a function of extracellular Ca^2+^ concentration. ACh was applied at near-EC50 concentrations (10 μM ACh for all α4β2, rat and chick α9α10 nAChRs and 100 μM ACh for all α7 and frog α9α10 nAChRs). Current amplitudes recorded at different Ca^2+^ concentrations in each oocyte were normalized to the response obtained at 1.8 mM Ca^2+^ in the same oocyte. V_h_: -90 mV. Bars represent mean ± S.E.M., open circles represent individual oocytes (n = 4-12). \**p*<0.05, \*\**p*<0.01, \*\*\**p*<0.005, \*\*\*\**p*<0.0001, Paired *t*-test (rat and frog α4β2 nAChRs and all α7 and α9α10 nAChRs) or Wilcoxon matched pair test (chick α4β2 nAChR) – comparing 0.5 mM Ca^2+^ vs 3 mM Ca^2+^.

Calcium permeation through nAChRs holds great functional significance for the activation of calcium dependent conductances and intracellular signalling pathways (14). Receptors containing the α7 subunit have high calcium permeability (40) whereas receptors containing α4β2 subunits have a lower contribution of calcium to the total current (28, 41). Amniote inner ear hair cell α9α10 nAChRs show differences in the extent of calcium permeability (23, 28). In order to perform a comparative analysis of the extent of the calcium component of ACh responses, we studied the differential activation of the oocyte’s endogenous calcium-activated chloride current (*I*Cl_Ca_). In oocytes expressing a recombinant receptor with high calcium permeability, the *I*Cl_Ca_ is strongly activated upon ACh application (42). Incubation of oocytes with the membrane-permeant fast Ca^2+^ chelator BAPTA-AM subsequently abolishes the Cl^-^ component of the total measured current (43). Fig. 6 shows representative responses to ACh before and after a 3-hour incubation with BAPTA-AM for α4β2, α7 and α9α10 nAChRs from rat, chicken and frog. Whereas ACh-evoked currents were only slightly affected in all α4β2 nAChRs denoting no major calcium influx (70-80% of current remaining after BAPTA incubation, Fig. 6, left panel), all α7 receptors showed a strong reduction of the ACh response after BAPTA incubation (30-40% of current remaining after BAPTA), indicating significant calcium permeation (Fig. 6, middle panel). Thus, no inter-species differences in the proportion of calcium current for both the low calcium permeant α4β2 and the highly calcium permeant α7 neuronal receptors were observed (Table S8). Conversely, and as previously reported (23, 28), we observed a marked difference in the extent of calcium current between the rat and chicken α9α10 receptors (p = 0.0299 – Fig. 6, right panels and Table S8). Moreover, the percentage of remaining response after BAPTA-AM incubation for the frog α9α10 receptor (Fig. 6, right panels) was similar to that of the rat receptor (p = 0.9999 – Table S8) and significantly different from that of the chicken α9α10 nAChR (p = 0.0013 – Table S8).

**Figure 6.**
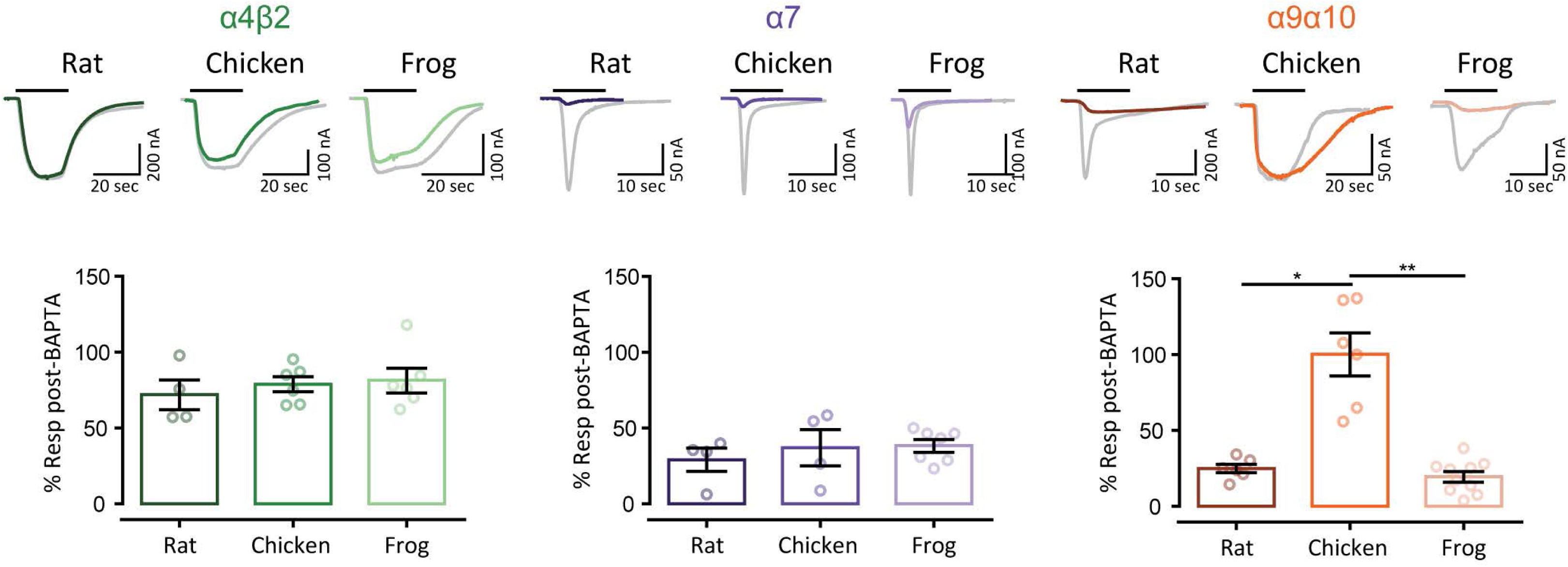
Unlike neuronal nAChRs, α9α10 nAChRs exhibit differential Ca^2+^ contribution to the total inward current. *Top panels*. Representative responses to near-EC_50_ concentration of ACh (10 μM ACh for all α4β2 and amniotes α9α10 nAChRs and 100 μM ACh for all α7 and frog α9α10 nAChRs) in oocytes expressing α4β2, α7 and α9α10 nAChRs, before (grey traces) and after (colour traces) a 3 hour incubation with BAPTA-AM (V_h_= −70 mV). *Bottom panels*. Percentage of the initial response remaining after BAPTA incubation. Bars represent mean ± S.E.M., open circles represent individual oocytes (n = 4-10). \*\*\*\**p*<0.0001, One-Way ANOVA followed by Dunn’s test (α4β2 and α7 nAChRs) or Kruskal-Wallis followed by Holm Sidak’s test (α9α10 nAChRs).

Neuronal nAChRs are characterised by a strong inward rectification, with negligible current at depolarized potentials (40, 41). This is proposed to be a relevant feature for their roles as pre-synaptic modulators of neuronal transmission (41). On the other hand, amniote α9α10 nAChRs exhibit a peculiar current-voltage (I-V) relationship due to a considerable outward current at positive potentials (19, 28). In order to perform a comparative analysis of the rectification profiles of neuronal and hair cell nAChRs we obtained I-V curves and determined the ratio of current elicited at +40 mV to that at -90 mV (*I*_+40_/*I*_-90_). All neuronal nAChRs exhibited I-V curves with marked inward rectification with no significant inter-species differences for either α4β2 or α7 tetrapod receptors, presenting *I*_+40_/*I*_-90_ values below 1 (Fig. 7, left and middle panels and Table S8). On the contrary, each of the hair cell α9α10 nAChRs analysed presented a unique I-V profile. As previously reported, rat α9α10 receptors showed significant outward current at depolarised potentials and greater inward current at hyperpolarised potentials with a *I*_+40_/*I*_-90_ ratio close to 1 (Fig. 7, right panels and (19)). The chicken α9α10 receptor showed outward current similar to that of the rat α9α10 receptor, but the inward current was smaller (Fig. 7, top right panel), resulting in a significantly higher *I*_+40_/*I*_-90_ ratio (p = 0.0229 – Table S8). Surprisingly, the frog α9α10 receptor showed an I-V profile similar to that of neuronal nAChRs, with strong inward rectification, almost no outward current at depolarised potentials (Fig. 7, right panels) and a *I*_+40_/*I*_-90_ below 1, significantly different to that obtained for chick (p = 0.0406 – Table S8) and rat (p < 0.0001 – Table S9) α9α10 receptors.

**Figure 7.**
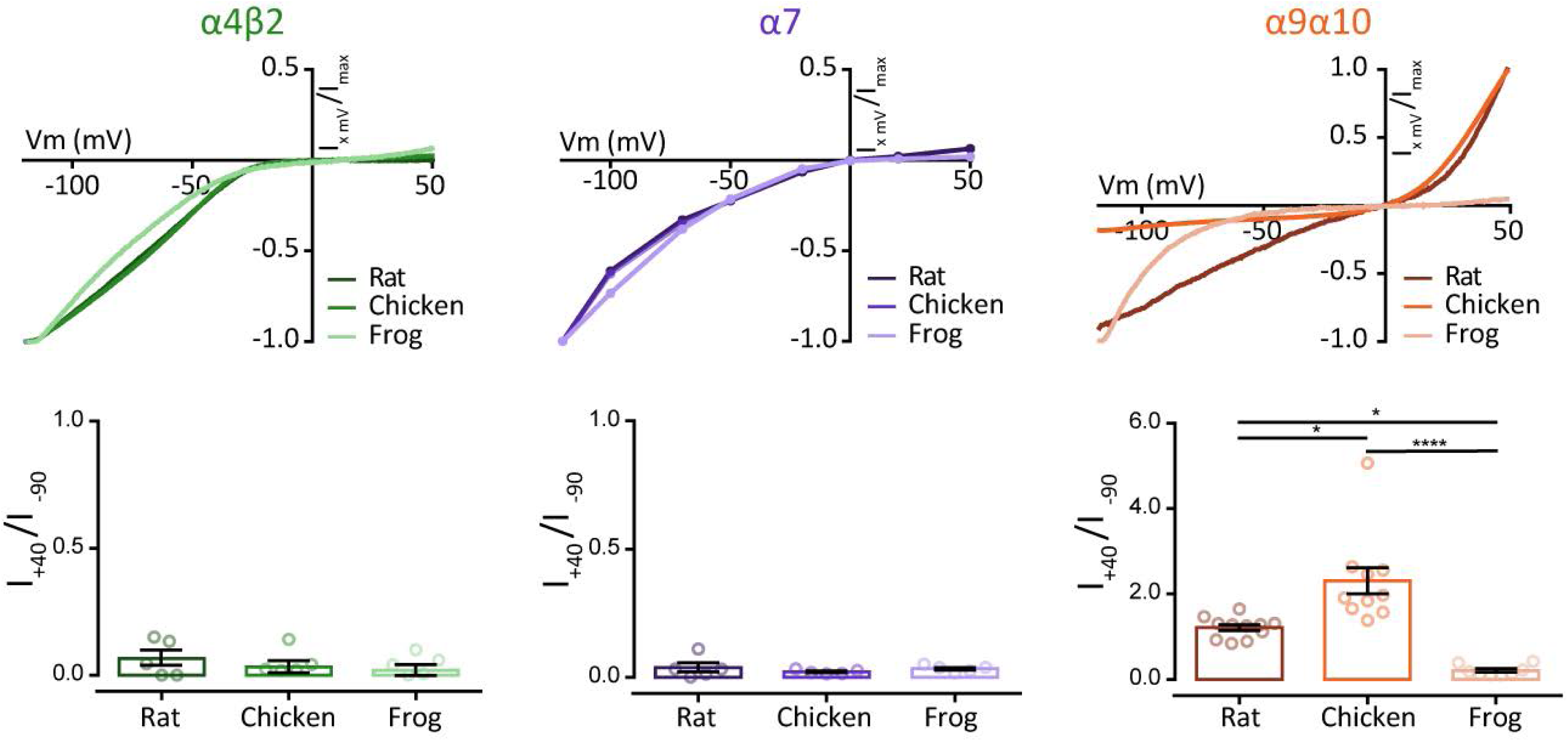
Hair cell, but not neuronal, nAChRs show differential current-voltage relationships across species. *Top panels*. Representative I–V curves obtained by the application of voltage ramps (-120 to +50mV, 2 seconds) at the plateau response to 3 μM ACh for α4β2 and α9α10 or by the application of 100 μM ACh at different holding potentials for α7 nAChRs. Values were normalized to the maximal agonist response obtained for each receptor. *Bottom panels*. Ratio of current amplitude at +40 mV relative to -90 mV for each oocyte. Bars represent mean ± S.E.M., open circles represent individual oocytes (n = 5-11). \**p*<0.05 \*\*\*\**p*< 0.0001, One-Way ANOVA followed by Dunn’s test (α4β2 and α7 nAChRs) or Kruskal-Wallis followed by Holm Sidak’s test (α9α10 nAChRs).

### Comparative functional analysis of neuronal and hair cell nAChRs shows distinct evolutionary trajectories

Altogether, the characterisation of individual functional properties of tetrapod nAChRs showed a stark contrast between neuronal and hair cell nAChRs. In order to concomitantly analyse the diversification, or conservation of receptor function, we performed principal component analysis (PCA) on all the functional variables measured on α4β2, α7 and α9α10 receptors from the three species (Table S10). The first two principal components accounted for 82% of the variability (Fig. 8). Moreover, the distribution of receptors in PCA space reflected their overall functional differences and similarities. Both neuronal α4β2 and α7 receptors occupied distinct regions, more distant in PC1 than in PC2 denoting that these receptors differ more on ACh apparent affinity, desensitisation and calcium permeability than they do on rectification and calcium modulation (Fig. 8 - inset). Also, α4β2 and α7 receptors from the different species were located very close together, reflecting the lack of inter-species differences in functional properties. In contrast, the hair cell α9α10 receptors from the different species were distant from each other in PCA space, denoting their extensive functional divergence (Fig. 8). Interestingly, the frog α9α10 nAChR was closer to the α7 receptors than to its amniote counterparts, highlighting the overall functional similarity between the amphibian α9α10 and α7 nAChRs.

**Figure 8.**
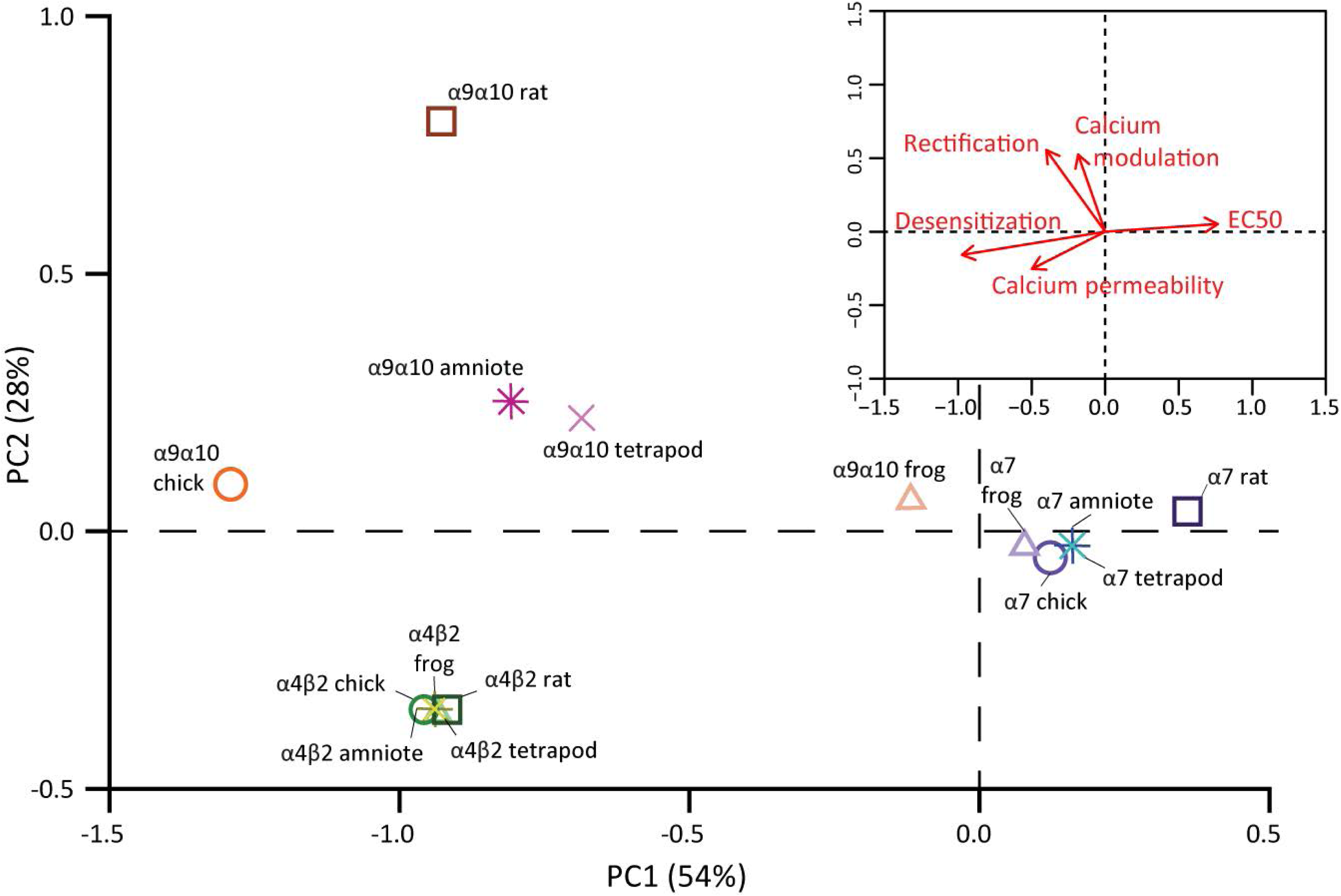
Hair cell nAChRs show great functional divergence, while functional properties of neuronal nAChRs are conserved. PCA was conducted using the experimentally determined biophysical properties (Table S8). Square symbols represent rat nAChRs, circles represent chick nAChRs and triangles represent frog nAChRs, α4β2 nAChRs are shown in shades of green, α7 nAChRs in shades of purple and α9α10 nAChRs in shades of orange. The projected locations of inferred functional states are shown for amniote (stars) and tetrapod (crosses) ancestral receptors and coloured in yellow (α4β2), blue (α7) or pink (α9α10). *Inset*. Biplot of the relative contribution of the five biophysical properties to PC1 and PC2.

Amino acid sequence phylogenies, co-expression/co-assembly patterns and functional experiments support the hypothesis of differential evolutionary trajectories for neuronal and hair cell receptors. Further insight could be attained by the evaluation of the receptors present in the last common ancestor of rat and chicken (amniote ancestor) and of rat, chicken and frog (tetrapod ancestor). To tackle this, we inferred the character state for the functional properties of the α4β2, α7 and α9α10 ancestral receptors (see SI methods). We then projected this predicted functional states onto the functional PCA space of the extant receptors (Fig. 8). As expected, the predicted ancestral α4β2 and α7 amniote and tetrapod receptors were located close to their extant counterparts. In contrast, the predicted ancestral states of the α9α10 receptors were localised halfway between the frog α9α10 and the amniote chicken and rat α9α10 extant receptors, potentially reflecting the functional evolutionary trajectory of the hair cell receptor. Thus, the ancestral tetrapod α9α10 receptor may have functionally resembled the extant frog α9α10 receptor and neuronal α7 receptors, mainly on properties with heavier loading on PC2 (i.e., rectification and calcium permeability). Moreover, the ancestral amniote α9α10 receptor may have had functional properties that more closely resembled those of neuronal α4β2 receptors, in particular for properties with heavier loading on PC1 (i.e., ACh apparent affinity, desensitization and calcium modulation). In summary, the data portrayed in Fig. 8 describes a plausible scenario for the functional evolutionary trajectories undertaken by two neuronal and the hair cell nAChRs within the tetrapod lineage.

## Discussion

The expansion of ion channel families on the vertebrate stem branch was mostly driven by two rounds of whole genome duplications (44). This has played a crucial role in the evolution of nervous systems and provided raw material that enabled the diversification of cell types (45) resulting in the complexity reached by vertebrate brains (e.g.(46)). Among the different ion channel families, Cys-loop receptors are within those that underwent the greatest expansion (45). The entire extant complement of nAChRs, which includes 17 different subunits (α1-α10, β1-β4, δ, ε and γ), was already present in the last common ancestor of all vertebrates (10). With the exception of some fish species that acquired α11, β1.2 and β5 subunits, for which no expression or functional data has yet been reported (10), and the loss of the α7-like α8 subunits in mammals, the complement of original vertebrate nAChR subunits has been remarkably conserved. Moreover, nAChRs are unique in that subgroups of the family have distinct roles in synaptic transmission in non-overlapping domains, either in the nervous system, inner ear hair cells or the neuromuscular junction (13, 14, 18). In this work, we performed in-depth analyses of coding sequence molecular evolution, subunit co-expression patterns and comparative functional properties of neuronal and hair cell receptors to explore the potential impact of these segregation of nAChRs subgroups on their respective evolutionary histories. We found that neuronal subunits showed high degree of coding sequence conservation, coupled to greater co-expression variability and conservation of functional properties across tetrapod clades. In contrast, the hair cell α9α10 nAChR showed greater sequence divergence, a highly restricted co-expression pattern and a great degree of variability of functional properties across species. These results indicate that along the tetrapod lineage neuronal and hair cell nAChRs underwent alternative evolutionary trajectories.

### Functional conservation of neuronal nAChRs

The observation that the biophysical functional properties of neuronal α4β2 and α7 nAChRs were conserved in the three tetrapod species analysed, relates to their high degree of amino acid sequence conservation (Fig. 1 and Tables S2-S3). Cholinergic innervation is pervasive, with almost every area of the brain being influenced by nicotinic signalling (14). Moreover, the expression of neuronal nAChR subunits is widespread in the central and peripheral nervous system (Fig. 2), where they assemble in multiple combinations (Table S6). Thus, randomly acquired changes in the coding sequence of a given subunit might be deleterious for receptor function in a multitude of heteropentameric assemblies present in diverse neuronal types. Such changes would therefore be under strong negative selection pressure. This is in agreement with the low degree of divergence observed for neuronal subunits and the absence of signatures of positive selection in the protein-coding sequences of α4, β2 and α7 subunits (28) and other brain expressed genes (47). However, neurons could alternatively resort to re-shuffling of the complement of nAChR subunits being expressed to attain functional divergence. Therefore, and as for most brain expressed genes (47, 48), random changes in non-coding regions that lead to differential expression patterns across brain areas or species may have played a substantial role in delineating the evolutionary trajectories of neuronal nAChRs.

### Differential co-expression patterns of neuronal and hair cell nAChRs

Our meta-analysis of expression patterns across the mouse brain highlights numerous instances of potential functional variability and diversification, even between closely related neuronal types (Fig. 2, S4-S6). For instance, cholinergic input controls dopaminergic neuron firing patterns in the midbrain (49). Here co-expression of nAChR subunits greatly differs across the four subtypes of dopaminergic neurons in the ventral tegmental area (VTA) ((50) and Fig. 2), indicating that they may express differential levels of functionally distinct α4β2 containing (α4β2*) receptors. Inclusion of the α5 subunit can alter α4β2* receptor properties substantially, increasing ACh sensitivity, desensitization kinetics and Ca^2+^ permeability (51–53). In addition, incorporation of the β3 subunit to the α4β2* receptor also increases ACh sensitivity, without significantly affecting Ca^2+^ permeability (51, 53). Moreover, VTA dopaminergic neurons also showed expression of the α6 and α3 subunits, both of which can co-assemble with α4, α5 and/or β2 subunits, greatly enhancing the complexity of individual nAChRs-mediated responses of VTA dopaminergic neurons to modulatory cholinergic input. Another interesting example is provided by layer VI cortical pyramidal neurons, whose activity is modulated by a dense cholinergic innervation from the basal forebrain. Here, ACh elicits robust excitatory responses acting on α4β2* nAChRs, with layer VI being one of the few cortical areas that express the accessory α5 nAChR subunit (54). Cortical neurons that project to both the ventral posteromedial nucleus (VPM) and the posteromedial complex of the thalamus express significantly higher levels of the α5 subunit than neurons projecting to the VPM only (Fig. S6A). This suggests VPM-only projecting neurons could have a lower density of α4α5β2 compared to α4β2 nAChRs, potentially contributing to differential cholinergic modulation of these subtypes of layer VI neurons, that also show differences in excitability (55).

Hair cells of the inner ear express high levels of α9 and α10 subunits, along with a number of neuronal (β2 and β4) and muscle (α1, β1, γ/ε) subunits (Fig. 2 and (20, 56)). α9α10 nAChRs mediate fast efferent neurotransmission to cochlear and vestibular hair cells in the inner ear (13) and are characterised by unique pharmacological and biophysical properties (19, 22, 39, 57). Most notably, nicotine, the diagnostic agonist of the nAChRs family, does not act as an agonist of α9α10 receptors, but as a competitive antagonist (19). In inner ear hair cells, no responses to nicotine application are detected (20, 58), indicating that functional muscle or neuronal nAChRs are not present at the plasma membrane. The presence of neuronal and muscle subunits mRNA may result from redundant or residual transcriptional regulation mechanisms. Moreover, similar “leaky” expression of muscle subunits was detected in a number of neuronal types (Fig. 2).

Expression of α9 and α10 subunit is restricted to inner ear hair cells, with a few interesting exceptions. In the inner ear, spiral ganglion neurons (SGNs) provide afferent innervation to cochlear hair cells and express a range of neuronal nAChR subunits (Fig. 2 and (59, 60)). Interestingly, low levels of α9 and α10 subunits are present in SGNs (Fig. 2 and (61)) with similarly low co-expression detected by two independent single-cell RNA sequencing studies (62, 63). If this low level of α9 and α10 mRNA proves to be more than “transcriptional noise”, then SGNs may be unique among neurons in expressing the hair cell α9α10 receptor. This might be related to the shared developmental origin of SGNs and hair cells at the otic placode (64, 65) and could open the possibility that in addition to neuronal nAChRs, which are thought to partly mediate the nicotinic effect of lateral olivocochlear terminals on afferent dendrites (66), α9α10 nAChRs may also play a role. Finally, in dorsal root ganglia neurons, α9 expression was not detected, while α10 is present at very low levels in only a few subtypes (Fig. 2 and (67)). These observations support those reported by qPCR and functional assays (68) and provide further evidence that the participation of α9* nAChRs in pain processes is not due to its presence in dorsal root ganglia neurons (68, 69).

Of note, α9 and α10, together with other nAChR subunits, are expressed in other peripheral, non-neuronal, tissues (17, 70). A plausible autocrine/paracrine effect of ACh in these cells can be served by a multiple and redundant battery of nAChRs that might play a signalling function in these peripheral tissues (71, 72). Due to the redundancy in pathways for ACh signalling, it is unlikely that the function of α9α10 nAChRs in these peripheral tissues provided the selection forces that shaped the accumulation of non-synonymous changes on this receptor.

### The α9α10 nAChR and the evolution of the efferent olivocochlear system

The observations that α9 and α10 genes are only co-ordinately transcribed in inner ear hair cells, together with their ability to only form functional heteromeric receptors when co-assembled with each other but not with other nAChR subunits (19, 20, 22), support our hypothesis that evolutionary changes in the hair cell receptors may have been focused at the coding sequence. Accordingly, vertebrate α9 and α10 subunits exhibit significant sequence divergence (Table S3 and Fig. 1), with mammalian α10 subunits showing a higher than expected accumulation of non-synonymous substitutions that were positively selected (26, 28). In addition, both α9 and a10 subunits show a high number of clade-specific (mammalian vs sauropsid) functionally relevant aminoacid changes (Fig. 1, Table S3 and (23)). Consequently, the biophysical properties of α9α10 receptors drastically changed across vertebrate species (Fig. 8 and (23, 28)). Since the primary function of the α9α10 receptor is at the postsynaptic side of the olivocochlear synapse, it can be hypothesised that clade specific differences in efferent modulation of hair cell activity could have shaped the functional properties of α9α10 receptors. Upon the transition to land, the hearing organs of tetrapods underwent parallel evolutionary processes, mainly due to the independent emergence of the tympanic middle ear, at least five times, in separate groups of amniotes (73). This was followed by the independent elongation of the auditory sensory epithelia that extended the hearing range to higher frequencies and the elaboration of passive and active sound amplification mechanisms that lead to the fine tuning of sound detection (73, 74). More importantly, mammals and sauropsids underwent a parallel diversification of hair cell types, segregating, partially in birds but completely in mammals, the phonoreception and sound amplification functions (75). Efferent innervation to hair cells, mediated by α9α10 nAChRs, is an ancestral feature common to all vertebrate species (76). In the auditory epithelia it modulates sound amplification and followed the hair cell diversification pattern: in birds it mainly targets short hair cells, while in mammals it targets outer hair cells (75). The latter developed a clade-specific sound amplification mechanism driven by the motor protein prestin and termed somatic electromotility (74). Prestin, together with βV giant spectrin, a major component of the outer hair cells’ cortical cytoskeleton which is necessary for electromotility, show signatures of positive selection in the mammalian clade that may relate to the acquisition of somatic electromotility (26, 77). Thus, the mammalian clade-specific evolutionary processes observed in both the α9 and α10 subunits (23, 26, 28) may be related to overall changes in the efferent olivovochlear systems of this clade that is tasked with the modulation of prestin-driven somatic electromotility. A recent high throughput evolutionary analysis identified 167 inner ear expressed genes with signatures of positive selection in the mammalian lineage (78). These inner ear genes, including those encoding the α9 and α10 nAChR subunits, can be considered as hotspots for evolutionary innovation in the auditory system across species.

Such a scenario provides a context for evaluating the relationship between evolutionary trajectories and the functional role of α9α10 receptors. In mammals, the high calcium influx through α9α10 receptors activates large conductance, voltage and low-calcium-sensitive BK potassium channels mediating hyperpolarization of outer hair cells in higher frequency regions of the cochlea (79). In contrast, in short hair cells from the chicken basilar papillae, hyperpolarization is served by the ACh-dependent activation of high calcium sensitive SK potassium channels (80, 81). Moreover, in contrast to adult mammalian hair cells where efferent fibres directly contact outer hair cells, but not the inner hair cells that release glutamate to activate afferent auditory fibres, efferent innervation in birds and amphibians co-exists with afferent innervation in the same hair cells. Calcium influx in these hair cells could therefore result in efferent-triggered, ACh-mediated release of glutamate to auditory afferents due to calcium spill over, bypassing sound mechanotransduction. Thus, limiting the extent of calcium influx through α9α10 nAChRs may be paramount to avoid confounding sensory inputs. In this hypothetical scenario, the low calcium permeability of the avian α9α10 nAChR or the very high desensitization kinetics of the amphibian α9α10 nAChR that restrict calcium load could be related to the aforementioned selection pressure.

### Subgroups of nAChRs and differential sources of functional divergence

Our observations on expression pattern, coding sequence and functional divergence support the notion that α9 and α10 are not a subtype of brain nicotinic subunit (for review see (25)), but form a group of their own, characterised by unique expression profile, pharmacological and biophysical properties (19, 22, 39, 57) and evolutionary history.

The contrasting evolutionary trajectories of neuronal and hair cell receptors, with functional variability stemming from combinatorial co-expression for the former and changes in coding sequence for the latter, support the notion of differential substrates for random change and ensuing functional divergence. For neuronal subunits, the source of random variability may have been rooted on changes in regulatory sequences. In contrast, for the hair cell receptor, random changes in the coding sequence were fixed throughout the evolutionary history of the tetrapod lineage. Interestingly, muscle subunits showed relatively low levels of coding sequence conservation (Fig. 1) and, via combinatorial co-assembly, muscle cells can toggle between at least two receptor variants (18). This places muscle receptors in between the two extremes of hair cell (isolated) and neuronal (widespread) receptors. A comparative functional study of muscle receptors would further test our hypothesis, with the prediction that a modest level of functional divergence may be encountered, but outweighed by the functional differences between muscle receptor variants.

In summary, the present work provides evidence supporting different evolutionary trajectories for neuronal and hair cell nAChRs. These may have resulted from the differential substrates for random change that dominated evolutionary processes in each receptor subgroup: diversity of co-expression/co-assembly patterns for neuronal subunits, changes in coding sequence for hair cell subunits. It results salient that among ligand-gated ion channels these alternative evolutionary trajectories are a particular feature of the nAChRs family of Cys-loop receptors. Thus, this is not observed for ionotropic glutamate and GABA_A_ receptors, whose 18 (82) and 19 (83) member subunits, respectively, are all expressed in the central nervous system. Finally, the simultaneous analysing of coding sequences, expression patterns and protein functional properties generated new insights into the evolutionary history of gene paralogues, thus providing further context for the role of nAChRs in neuronal and hair cell synaptic transmission.

## Methods

All experimental protocols were carried out in accordance with the guides for the care and use of laboratory animals of the National Institutes of Health and the Institutional Animal Care and Use Committee of the Instituto de Investigaciones en Ingeniería Genética y Biología Molecular, “Dr. Héctor N. Torres”.

### Phylogenetic analysis of vertebrate nAChRs subunits

All sequences were downloaded from GenBank (www.ncbi.nlm.nih.gov/genbank), UCSC (http://genome.ucsc.edu/) and Ensembl (www.ensembl.org) databases. Sequence alignment was performed using *Clustal*W on the MEGΑ7 software (84). Phylogenetic trees were inferred using the minimum evolution method (85). A detailed description of the alignment procedures, sequence identity analysis and phylogeny constructions are available in SI Methods.

### Functional divergence analysis

The coefficient of type II functional divergence (θ_II_), its standard error and the site-specific posterior probabilities were calculated for each nicotinic subunit between the mammalian and sauropsid clades using DIVERGE 3.0 (31). A detailed description of the analysis strategy is available in SI methods.

### Analysis of nAChR subunit expression in single-cell RNAseq (scRNAseq) datasets

A meta-analysis of single cell gene expression data from 10 studies was performed to describe the expression patterns of nAChR subunits across cell types. Probability distributions of mean expression level for all nAChR subunit genes detected in all the cell types analysed were obtained using the *scde* package (32). We combined this with a comprehensive catalogue of experimentally validated subunit combinations and identified the subunit combinations that were present, in each cell type or absent all together. A detailed description of the datasets used and analysis strategy is available in SI methods.

### Expression of recombinant receptors in Xenopus laevis oocytes and electrophysiological recordings

The plasmids used for heterologous expression in *Xenopus laevis* oocytes are described in SI Methods. Electrophysiological recordings from *Xenopus laevis* oocytes were obtained as described previously (22, 57) and are described in more detail in SI Methods. A detailed description of the protocols used to evaluate nAChR functional properties and statistical analysis is available in SI methods.

### Principal component analysis of functional properties and inference of character state of functional properties of ancestral receptors

Principal Component Analysis was performed on the experimental values obtained for the functional properties of extant nAChRs implementing custom routines written in R v3.4.1 and run in RStudio software v1.0.153 (see SI methods). The pipeline for the inference of the ancestral character state of biophysical properties is described in detail in SI methods. Briefly, sequences corresponding to the amniote and tetrapod ancestors were reconstructed for the different subunits. Distances were then used to compute branch lengths on receptor trees. The latter, together with the biophysical properties of extant receptors were used to infer the properties of ancestral receptors using the APE package v5.2 (86). Finally, these were projected onto the PCA space of extant receptors.

## Acknowledgments

This work was supported by Agencia Nacional de Promoción Científicas y Técnicas, Argentina, the Scientific Grand Prize of the Fondation Pour l’Audition and NIH Grant R01 DC001508 (Paul Fuchs PI and ABE co-PI) to ABE.

## Supplementary methods

### Phylogenetic analysis of vertebrate nAChRs subunits

All sequences were downloaded from GenBank (www.ncbi.nlm.nih.gov/genbank), UCSC (http://genome.ucsc.edu/) and Ensembl (www.ensembl.org) databases. The signal peptides of all sequences were excluded from the analysis since they are not present in the mature functional protein. Accession numbers are listed in Table S1. All sequences were visually inspected, and missing and/or incorrect exons were obtained from the NCBI Genome Project traces database (http://blast.ncbi.nlm.nih.gov/Blast.cgi). Sequence alignment was performed using *Clustal*W on the MEGΑ7 software (www.megasoftware.net; (1)), with the following parameters: for pairwise alignments, gap opening penalty: 10, gap extension penalty: 0.1; for multiple alignments, gap opening penalty: 10, gap extension penalty: 0.2. Protein weight matrix: Gonnet. Residue specific and hydrophilic penalties: ON. Gap separation distance: 4. End gap separation: OFF and no negative matrix was used. The delay divergent cutoff was 30%. The full alignment for the nicotinic subunits from representative vertebrate species is available in Supplementary File 1 in fasta format.

The phylogenetic tree of all nAChR subunits was built using the MEGΑ7 software. Positions containing alignment gaps and missing data were eliminated only in pairwise sequence comparisons. The final dataset contained a total of 773 positions. The evolutionary distances (i.e., number of amino acid substitutions per site) were computed using the JTT matrix-based method (2). The neighbour-joining algorithm (3) was used to generate the initial tree and branch support was obtained by bootstrap test (1000 replicates) (4). The evolutionary history was inferred using the minimum evolution method (5). A first tree was generated with variation rate among sites modelled by a gamma distribution (Figure S1) and a second tree assuming uniform variation rates among sites (Figure S2). Overall, tree topology was similar between both methods.

Average percentage sequence identity was calculated for each subunit using the percentage of sequence identity between each pair of sequences (Supplementary File 2) from the same category for all sequences and for within or between mammalian and/or sauropsid sequences. For α10 subunits, the average percentage of sequence identity was also calculated for the non-mammalian paraphyletic group. Values obtained are summarised in Table S2.

The cumulative distribution of percentage sequence identity between pairs of sequences from all amniotes and within or between mammalian and/or sauropsid species was plotted for each subunit (Fig. S3).

### Functional divergence analysis

The DIVERGE 3.0 software was used to statistically test for functional diversification of nAChR subunits between the mammalian and sauropsid clades. DIVERGE predicts amino acid sites that may be involved in between-clade functional divergence against the background of neutral evolution (6). In particular, it estimates the type II functional divergence coefficient (θ_II_) that indicates site-specific evolutionary shifts in aminoacid biochemical state between clades and then uses a Bayesian approach to compute the posterior probability that each individual site contributes to the clade-specific functional diversification. Type II sites represent aminoacids that are highly conserved in each clade, but in a biochemically different state (i.e., positively charged in clade 1 and negatively charged in clade 2).

Multiple alignments of protein sequences for each individual subunit were generated using the MEGA7 software as described above. The highly variable amino-terminal signal peptides and intracellular domains were excluded from the analysis. Phylogenetic trees, with a topology corresponding to the species tree, were constructed for each nicotinic subunit by Maximum Likelihood and the JTT matrix-based method (2). The rates among sites were modelled as a gamma distribution. All positions with less than 95% site coverage were eliminated. α8, β1 and ε nAChR subunits were not included in the analysis due to lack of mammalian and/or sauropsid sequences to perform suitable comparisons.

Multiple alignments of protein sequences and their corresponding phylogenetic trees (Supplementary File 3) were used as input data for DIVERGE 3.0 type II functional divergence analysis (7), with default parameters. θ_II_ and site-specific posterior probabilities were calculated for each subunit. A θ_II_ value significantly greater than 0 (P < 0.05) indicates that residues conserved within each group have undergone radical changes in amino acid identity between groups (6). z-scores were used to test the significant difference of θ_II_ coefficients against the null hypothesis (θ_II_ = 0) that implies no sites are present in the protein that reflect between-clade functional divergence (Table S3). Site-specific posterior probabilities were computed for all sites along each subunit (Supplementary File 3). Sites with posterior probabilities greater than 0.65 for each subunit are highlighted in Figure 1B.

### Analysis of nAChR subunit expression in single-cell RNAseq (scRNAseq) datasets

Processed gene expression data tables were obtained from 10 scRNAseq studies that evaluated gene expression in retina (8) inner ear sensory epithelium (9, 10) and spiral ganglion (11), ventral midbrain (12), hippocampus (13), cortex (14), hypothalamus (15), visceral motor neurons (16) and dorsal root ganglia (17). Accession numbers, cell types inferred and number of cells analysed are summarised in Table S4. For all datasets, we used the published gene expression quantification and cell type labels. Each dataset was analysed separately. For the retina dataset we used the Smart-Seq2 sequencing data from Vsx2-GFP positive cells (8). For the gene expression quantification we only analysed four cell types that had a minimum number of cells in the dataset that allowed reliable fitting to the error models: BC1A, BC5A, BC6 and RBC. From the layer VI somatosensory cortex dataset (13) we used a subset of the expression matrix that corresponds to day 0 (i.e. control, undisturbed neurons) of their experimental manipulation. For the hypothalamic neurons dataset (15) we used a subset that contained only neurons from untreated (control) mice and only quantified gene expression on the 10 broad cell types identified. From the ventral midbrain dopaminergic neurons dataset (12) we used a subset comprising DAT1-Cre/tdTomato positive neurons from P28 mice. For the SGNs dataset we used a subset comprising Type I neurons from wild type mice (11). For the utricle hair cell datasets we used the normalised expression data of (10). For the cochlear hair cell data we used the normalised expression data from (9) and continued our analysis with only the 10 cochlear hair cells identified. For the visceral motor neurons dataset (16) we excluded the neurons that were “unclassified” from further analysis. For the dorsal root ganglia dataset (17) we used a subset containing only successfully classified neurons that were collected at room temperature. Inspection of all datasets for batch effects was performed using the *scater* package (version 1.10.1) (18). Single cell expression of nAChR subunits was initially evaluated on violin plots (Fig. S4). All data analysis was implemented in R (version 3.5.1) and Bioconductor (version 3.8) (http://www.bioconductor.org/), running on RStudio (version 1.1.456) (http://www.rstudio.com/).

The publicly available expression matrices for a number of the datasets contained raw counts (retina, hippocampus, hypothalamus, midbrain, visceral motor neurons). For each of these dataset individually, we performed a normalisation step using the *scran* package (version 1.10.2) (19) that computes pool-based size factors that are subsequently deconvolved to obtain cell-based size factors.

The normalised expression matrices and cell type information were used as input to quantify cell-type specific gene expression. Analysis was performed using the *scde* package (version 1.99.1) (20). We modelled the gene expression measurements from each individual cell as a weighted mixture of negative binomial and low-magnitude Poisson distributions. The former accounts for the correlated component in which a transcript is detected and quantified, while the latter accounts for drop-out events in which a transcript fails to amplify when present. The weighting of the two distributions is determined by the level of gene expression in the cell population (20). We then used these error models to estimate the likelihood (joint posterior probability) of a gene being expressed at any given average level in a given cell type (20). Probability distributions for all nAChR subunit genes detected in all the cell types analysed are shown in Fig S5. This whole transcriptome analysis provides accurate estimations of gene expression levels, thus allowing for the comparison of individual genes within a given cell type (i.e. the complement of nAChR subunits) or the analysis of expression level differences between cell types (i.e. change in expression level of nAChR subunits between neuronal subtypes). Inferred mean expression values are summarised in Table S5. We combined the information about the complement of nAChR subunits for each cell type with a comprehensive catalogue of experimentally validated subunit combinations (Table S6 and references therein). We identified the subunit combinations that were present, in each cell type, within a 10-fold, 10 to 100-fold or 100 to 1000 fold range of expression level or absent all together. Admittedly, this analysis approach overlooks the complexities of post-translational modifications, receptor assembly, role of chaperone proteins and transport to the plasma membrane. However, it provides conservative estimates of the maximum potential of combinatorial assembly of subunits and thus a maximum for the repertoire of nAChR assemblies that could be present at the cell membrane.

### Expression of recombinant receptors in *Xenopus laevis* oocytes and electrophysiological recordings

Rat and chick α9 and α10 cDNAs subcloned into pSGEM, a modified pGEMHE vector suitable for *Xenopus laevis* oocyte expression studies, were described previously (21–23). Rat α4, β2 and α7 subunit cDNAs subcloned into pBS SK(−) (Agilent Technologies, Santa Clara, CA) were kindly provided by Dr. Jim Boulter (University of California, Los Angeles, CA). Chicken α4 and β2 subunit cDNAs subcloned into pCI (Promega, Madison, WI) were kindly provided by Dr. Isabel Bermudez-Díaz (Oxford Brookes University, Oxford, UK). Chicken α7 subunit cDNA cloned into pMXT was kindly provided by Dr. Jon Lindstrom (University of Pennsylvania) and was then subcloned into pSGEM between HindIII and SalI sites. Frog α4, β2, α7, α9 and α10 subunits were cloned from whole brain *Xenopus tropicalis* cDNA. Total RNA was prepared from whole brains using the RNAqueous – Micro kit AM1931 (Ambion, Thermo Fisher Scientific, Boston, MA). First strand cDNA synthesis was performed using an oligodT and the ProtoScript Taq RT-PCR kit (New England Biolabs, Ipswich, MA). Second strand synthesis was performed with the NEBNext mRNA Second Strand Synthesis Module kit – E6111S (New England Biolabs, Ipswich, MA). Full length cDNAs for each subunit were PCR amplified (MultiGene 60 OptiMaxTM Thermal Cycler - Labnet International Inc. Edison, NJ) using specific primers (Table S7). PCR products were sequenced and subcloned into pSGEM between EcoRI and XhoI sites for α9, α7 and β2 nAChRs subunits, between HindIII and XhoI sites for the α10 subunit and between EcoRI and HindIII sites for the α4 subunit. All expression plasmids are readily available upon request.

Capped cRNAs were *in vitro* transcribed from linearized plasmid DNA templates using the RiboMAX^TM^ Large Scale RNA Production System-T7 (Promega, Madison, WI). The maintenance of *Xenopus laevis*, as well as the preparation and cRNA injection of stage V and VI oocytes, has been described in detail elsewhere (Katz et al., 2000). Briefly, oocytes were injected with 50 nl of RNase-free water containing 0.01–1.0 ng of cRNAs (at a 1 : 1 molar ratio for α9α10 and α4β2 receptors) and maintained in Barth’s solution (in mM): NaCl 88, Ca(NO_3_)_2_ 0.33, CaCl_2_ 0.41, KCl 1, MgSO_4_ 0.82, NaHCO_3_ 2.4, HEPES 10, at 18°C.

Electrophysiological recordings were performed 2 – 6 days after cRNA injection under two-electrode voltage clamp with an Oocyte Clamp OC-725B or C amplifier (Warner Instruments Corp., Hamden, CT). Recordings were filtered at a corner frequency of 10 Hz using a 900BT Tunable Active Filter (Frequency Devices Inc., Ottawa, IL). Data acquisition was performed using a Patch Panel PP-50 LAB/1 interface (Warner Instruments Corp., Hamden, CT) at a rate of 10 points per second. Both voltage and current electrodes were filled with 3M KCl and had resistances of ∼1MΩ. Data were analysed using Clampfit from the pClamp 6.1 software (Molecular Devices, Sunnyvale, CA). During electrophysiological recordings, oocytes were continuously superfused (10 ml min^-1^) with normal frog saline composed of (in mM): 115 NaCl, 2.5 KCl, 1.8 CaCl_2_ and 10 HEPES buffer, pH 7.2. In order to minimize the activation of the oocyte’s native Ca^2+^-sensitive chloride current (*I*Cl_Ca_) by inward Ca^2+^ current through the nAChRs, all experiments, unless otherwise stated, were carried out in oocytes incubated with the membrane permeant Ca^2+^ chelator 1,2-bis (2-aminophenoxy)ethane-N,N,N’,N’-tetraacetic acid-acetoxymethyl ester (BAPTA-AM; 100 μM) for 3 h prior to electrophysiological recordings. This treatment was previously shown to effectively chelate intracellular Ca^2+^ ions and, therefore, to impair the activation of the *I*Cl_Ca_ (24). All recordings were performed at -70 mV holding potential, unless otherwise stated.

### Biophysical properties of nAChRs

Concentration-response curves were obtained by measuring responses to increasing concentrations of ACh. Current amplitudes were normalized to the maximal agonist response in each oocyte. The mean and S.E.M. values of the responses are represented. Agonist concentration-response curves were iteratively fitted, using Prism 6 software (GraphPad Software Inc., La Jolla, CA), with the equation: I/Imax = AnH / (AnH + EC_50_nH), where I is the peak inward current evoked by the agonist at concentration AnH; Imax is current evoked by the concentration of agonist eliciting a maximal response; EC_50_ is the concentration of agonist inducing half-maximal current response and nH is the Hill coefficient.

Desensitisation of ACh evoked currents was evaluated via prolonged agonist applications. The percentage of current remaining 5 seconds (for α7 nAChRs) or 20 seconds (for α4β2 and α9α10 nAChRs) after the peak of the response was determined for each oocyte.

The effects of extracellular Ca^2+^ on the ionic currents through nAChRs were studied by measuring the amplitudes of the responses to a near-EC_50_ concentration of ACh (10 μM for all α4β2 and amniote α9α10 nAChRs, and 100 μM for all α7 and frog α9α10 nAChRs) on extracellular Ca^2+^ ranging from nominally 0 to 3 mM at a holding potential of -90 mV (25). These experiments were carried out in oocytes injected with 7.5 ng of an oligonucleotide (5’-GCTTTAGTAATTCCCATCCTGCCATGTTTC-3’) antisense to connexinC38 mRNA (26, 27) to minimise the activation of the oocyte’s nonselective inward current through a hemigap junction channel that results from the reduction of external divalent cation concentration. Current amplitudes at each Ca^2+^ concentration were normalized to that obtained in the same oocyte at a 1.8 mM Ca^2+^.

I–V relationships were obtained by applying 2 seconds voltage ramps from -120 to +50 mV from a holding potential of -70 mV, at the plateau response to 3 μM ACh. Leakage correction was performed by digital subtraction of the I–V curve obtained by the same voltage ramp protocol prior to the application of ACh. Generation of voltage protocols and data acquisition were performed using a Patch Panel PP-50 LAB/1 interface (Warner Instruments Corp., Hamden, CT) at a rate of 10 points per second and the pClamp 7.0 software (Axon Instruments Corp., Union City, CA). Current values were normalized to the maximum amplitude value obtained for each oocyte. The fast desensitising α7 receptors had negligible plateau currents. For these receptors, responses to 100 μM ACh were obtained at different holding potentials and normalised to the amplitude response at -120 mV in the same oocyte.

Table S8 summarises the biophysical properties and statistical comparisons from rat, chicken and frog α4β2, α7 and α9α10 receptors.

### Statistical analysis

Shapiro-Wilks normality test was conducted using custom routines written in R v3.4.1 (R Development Core Team, 2008), through RStudio software v1.0.153. Statistical significance was determined using either parametric paired *t*-test or One-way ANOVA followed by Holm-Sidak’s test, or nonparametric Wilcoxon or Kruskal-Wallis tests followed by Dunn’s tests conducted using Prism 6 software (GraphPad Software Inc., La Jolla, CA). A p< 0.05 was considered significant.

### Reagents

All drugs were obtained from Sigma-Aldrich (Buenos Aires, Argentina). ACh chloride was dissolved in distilled water as 100 mM stocks and stored aliquoted at -20°C. BAPTA-AM was stored at -20°C as aliquots of a 100 mM stock solution in dimethylsulfoxide, thawed and diluted into Barth’s solution shortly before incubation of the oocytes. ACh solutions in Ringer’s saline were freshly prepared immediately before application.

### Principal component analysis of functional properties

Principal Component Analysis was performed on the experimental values obtained for the functional properties of extant nAChRs implementing custom routines written in R v3.4.1 and run in RStudio software v1.0.153. Each of the experimental variables was normalized to the maximum value recorded to allow for equal weighting of the properties (Table S9). The loadings of the empirical variables on the five principal components (PC) generated are shown in Table S10, alongside the proportion of the total variability explained by each component. The loadings of each biophysical property on each principal component are also shown on the vectors biplot (Figure 8 – inset).

### Inference of character state of functional properties of ancestral receptors

The pipeline followed to infer the ancestral character state of biophysical properties of nAChRs is schematized on Fig. S7. Briefly, we first reconstructed the ancestral tetrapods and amniote DNA sequences of the α4, α7, α9, α10 and β2 nAChRs subunits (Supplementary File 4). For that purpose, we used the same orthologue sequences that were used to construct the phylogenetic tree of Figure 1, together with a species tree with no branch lengths obtained from Ensembl (https://www.ensembl.org/info/about/speciestree.html). Inferred ancestral DNA sequences for the amniote and tetrapod nodes (Supplementary File 4) were obtained, for all three codon positions, by the maximum likelihood method (28) under the Tamura-Nei model (29), on the MEGΑ7 software (1). The initial tree corresponds to the provided Species Tree with very strong restriction to branch swap. The rates among sites were treated as a gamma distribution. All positions with less than 95% site coverage were eliminated.

Subsequently, multiple alignments including extant rat, chick and frog and ancestral amniote and tetrapod aminoacid sequences were performed using MEGΑ7 for each studied nAChR (Supplementary File 5). The sequence identity was used to assign branch length values to α7, α4β2 and α9α10 nAChRs trees, corresponding to 1-SeqID (Table S11). Theoretical concatemeric constructions were built for the heteromeric α9α10 nAChRs considering the descripted prevalent (α9)_2_(α10)_3_ stoichiometry (Plazas et al., 2005). For the α4β2 nAChRs, the high sensitivity (α4)_2_(β2)_3_ stoichiometry was used to generate the theoretical concatemeric receptor (Supplementary File 5).

The resulting trees (Fig. S8) were used, together with the biophysical properties experimentally determined for the extant receptors (Table S8) as input data for ancestral character inference. ACh sensitivity (EC_50_ values), desensitization rates (% of remaining *I* 20 sec after ACh peak), Ca^2+^ modulation (*I* elicited by ACh at Ca 0.5 mM/Ca 3 mM), Ca^2+^ permeability (% of remaining *I* after BAPTA treatment) and rectification profile (*I*_+40 mV_/*I*_-90 mV_) for the ancestral amniote and tetrapod receptors were inferred using the *ace* function from the *APE* package v5.2 (30) implemented in R v3.4.1 and RStudio v1.0.153. We used the Brownian motion model (31) (Schluter et al., 1997), where characters evolve following a random walk fitted by maximum likelihood (32) for the ancestral character estimations of continuous traits. Reconstructed ancestral states are shown in Fig. S9. Finally, using the loadings of the biophysical properties on PC1 and 2 (Table S10 and Figure 8 – inset), and the normalized *in silico* reconstructed biophysical properties inferred for the ancestral receptors we calculated their position on the bidimensional PCA space (Figure 8).

**Supplementary Figure 1.**
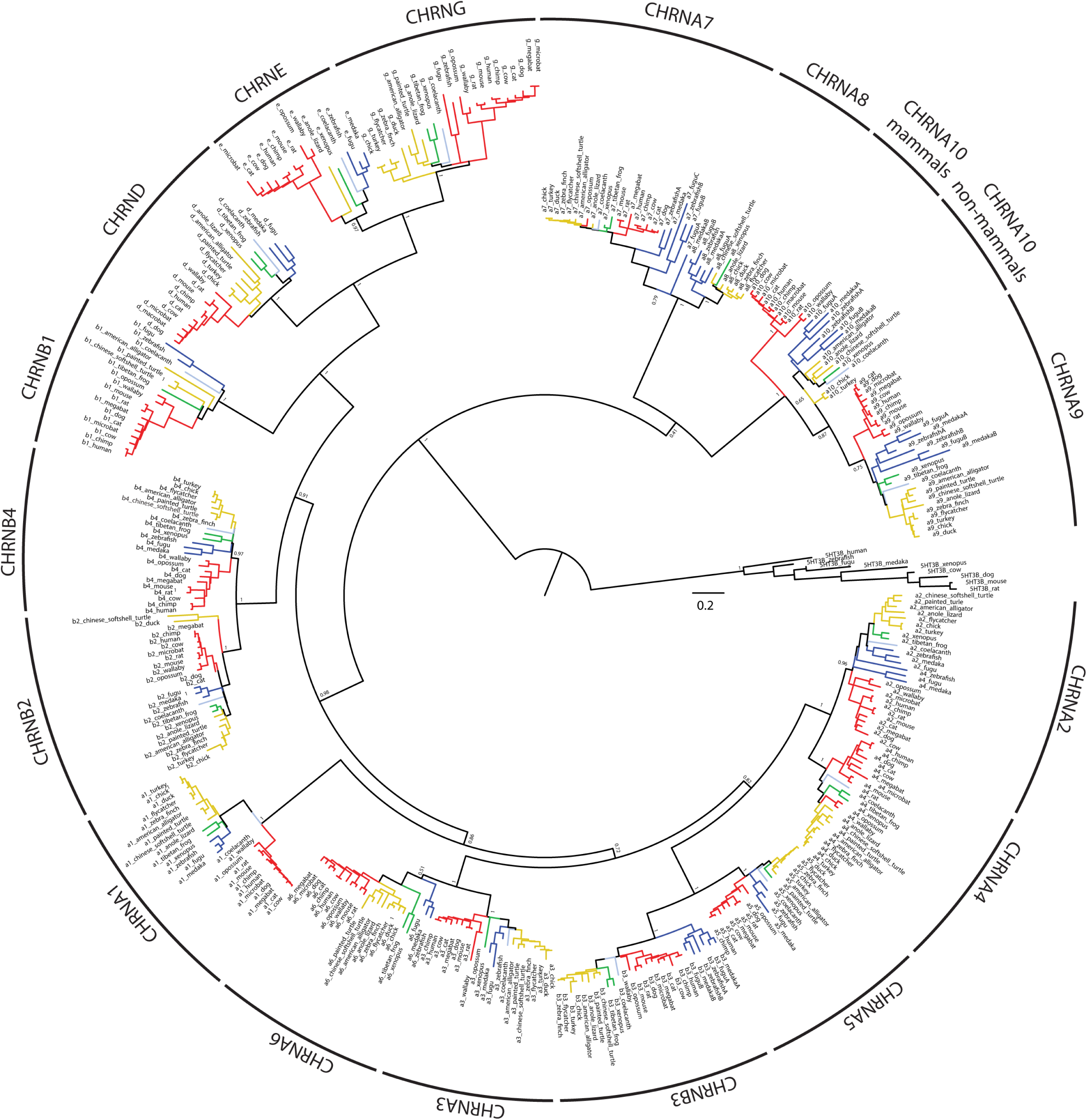
Complete minimum evolution phylogenetic tree corresponding to the collapsed tree shown in Figure 1, obtained with variation rates among sites modelled by a gamma distribution. Red branches, mammals; yellow branches, sauropsids; green branches, amphibians; blue branches, fish; light blue branches, coelacanth. The trees were built using minimum evolution method and pairwise deletion for missing sites. The optimal tree with a sum of branch length of 47.32565515 is shown. For clarity, the percentage of replicate trees in which the associated taxa clustered together in the bootstrap test (1,000 replicates) are shown only next to the branches that separate different subunits. The tree is drawn to scale, with branch lengths in the same units as those of the evolutionary distances used to infer the tree.

**Supplementary Figure 2.**
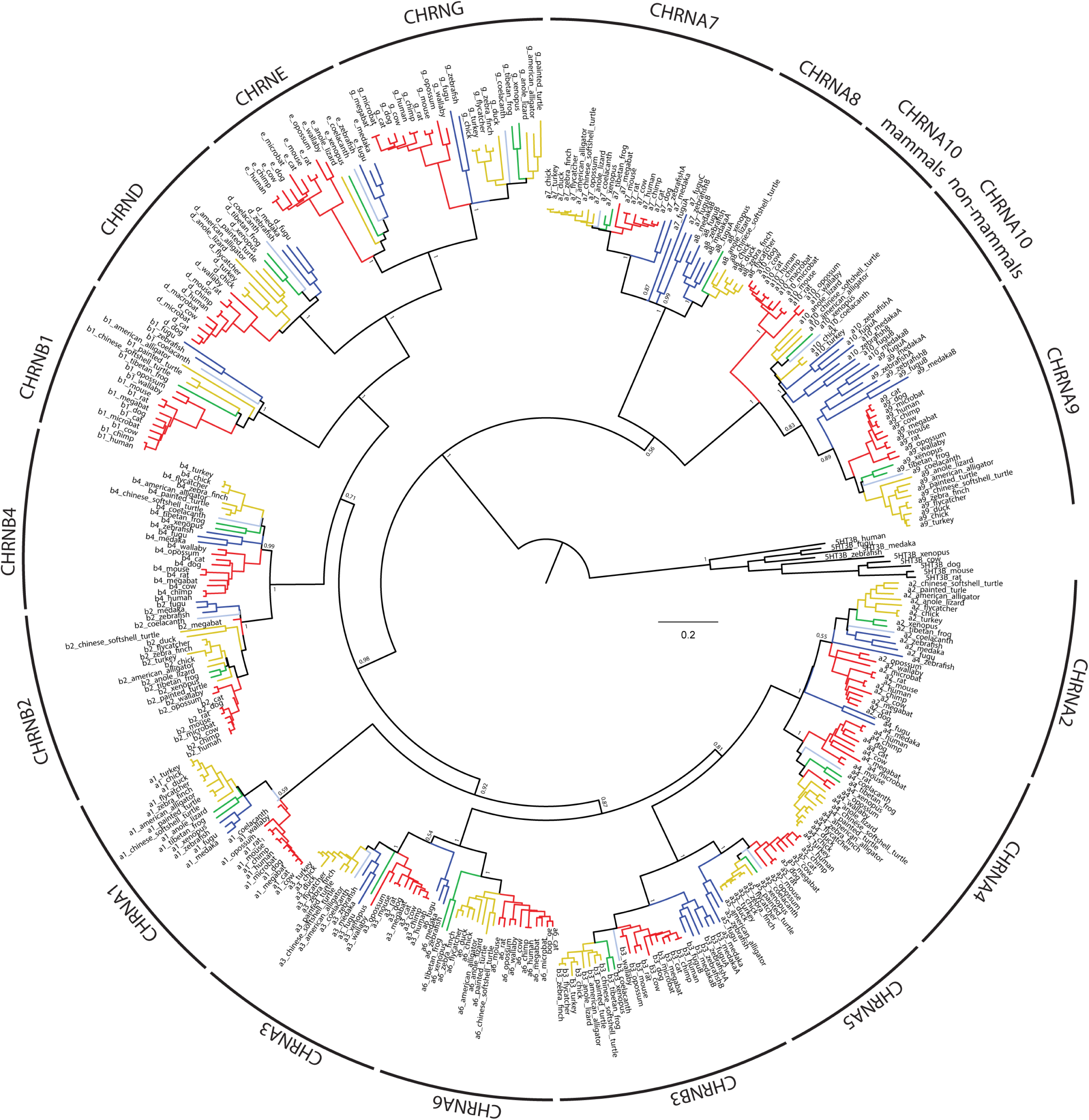
Complete minimum evolution phylogenetic tree obtained assuming uniform variation rates among branches. Red branches, mammals; yellow branches, sauropsids; green branches, amphibians; blue branches, fish; light blue branches, coelacanth. The trees were built using minimum evolution method and pairwise deletion for missing sites. The optimal tree with a sum of branch length of 37.49411418 is shown. For clarity, the percentage of replicate trees in which the associated taxa clustered together in the bootstrap test (1,000 replicates) are shown only next to the branches that separate different subunits. The tree is drawn to scale, with branch lengths in the same units as those of the evolutionary distances used to infer the tree.

**Supplementary Figure 3.**
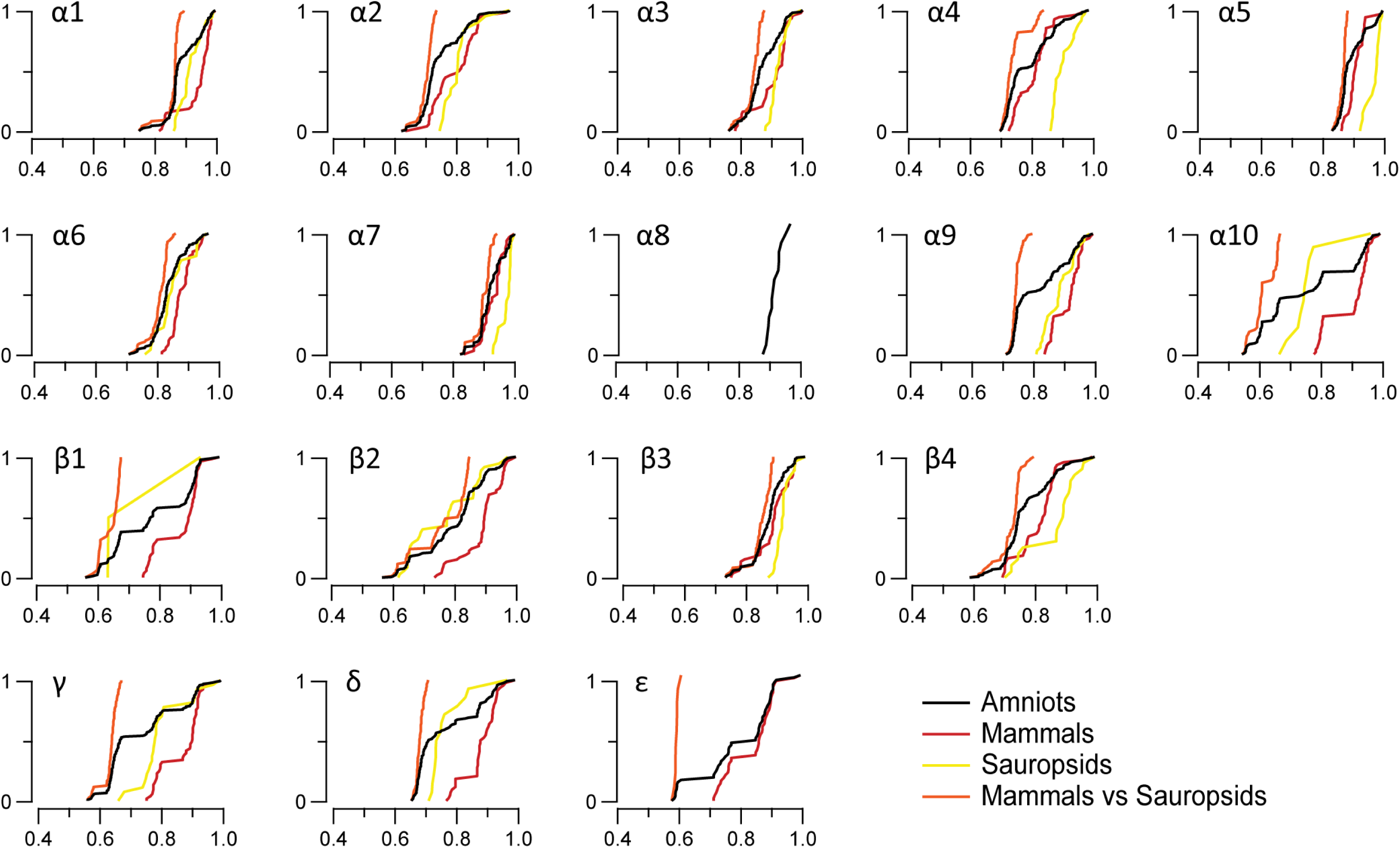
Cumulative distributions of sequence identity of individual nAChR subunits. Cumulative distributions were calculated from the sequence identity values obtained for each pair of subunits included in the following groups: only mammalian sequences (red), only sauropsid sequences (yellow), mammalian versus sauropsid sequences (orange) and all mammalian and sauropsid (amniote) sequences (grey).

**Supplementary Figure 4.**
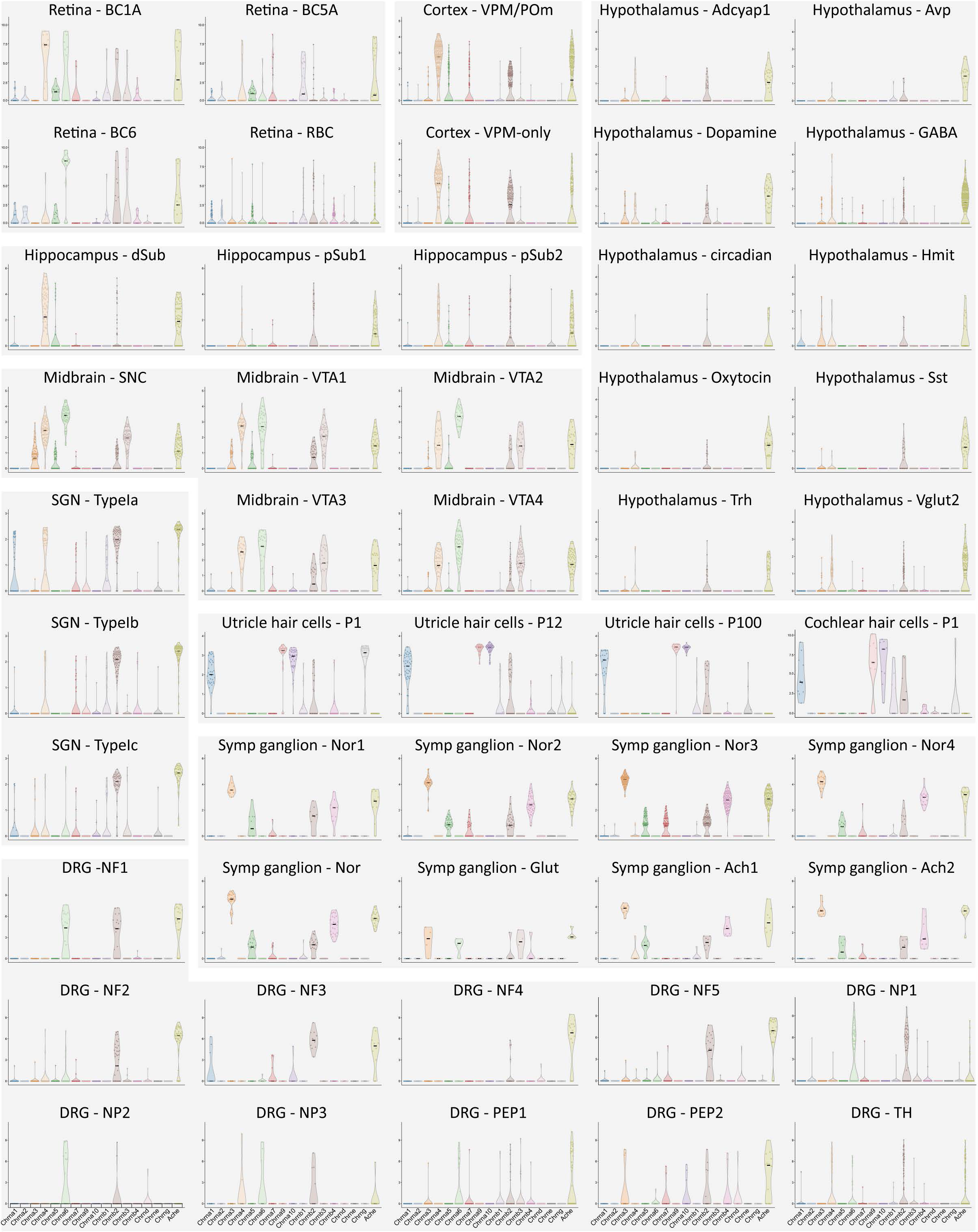
Violin plots showing expression level for all nAChR subunits and acetylcholinesterase (AChE) in the different cell types from the datasets analysed. Expression values are in log2 normalised counts (see methods for details).

**Supplementary Figure 5.**
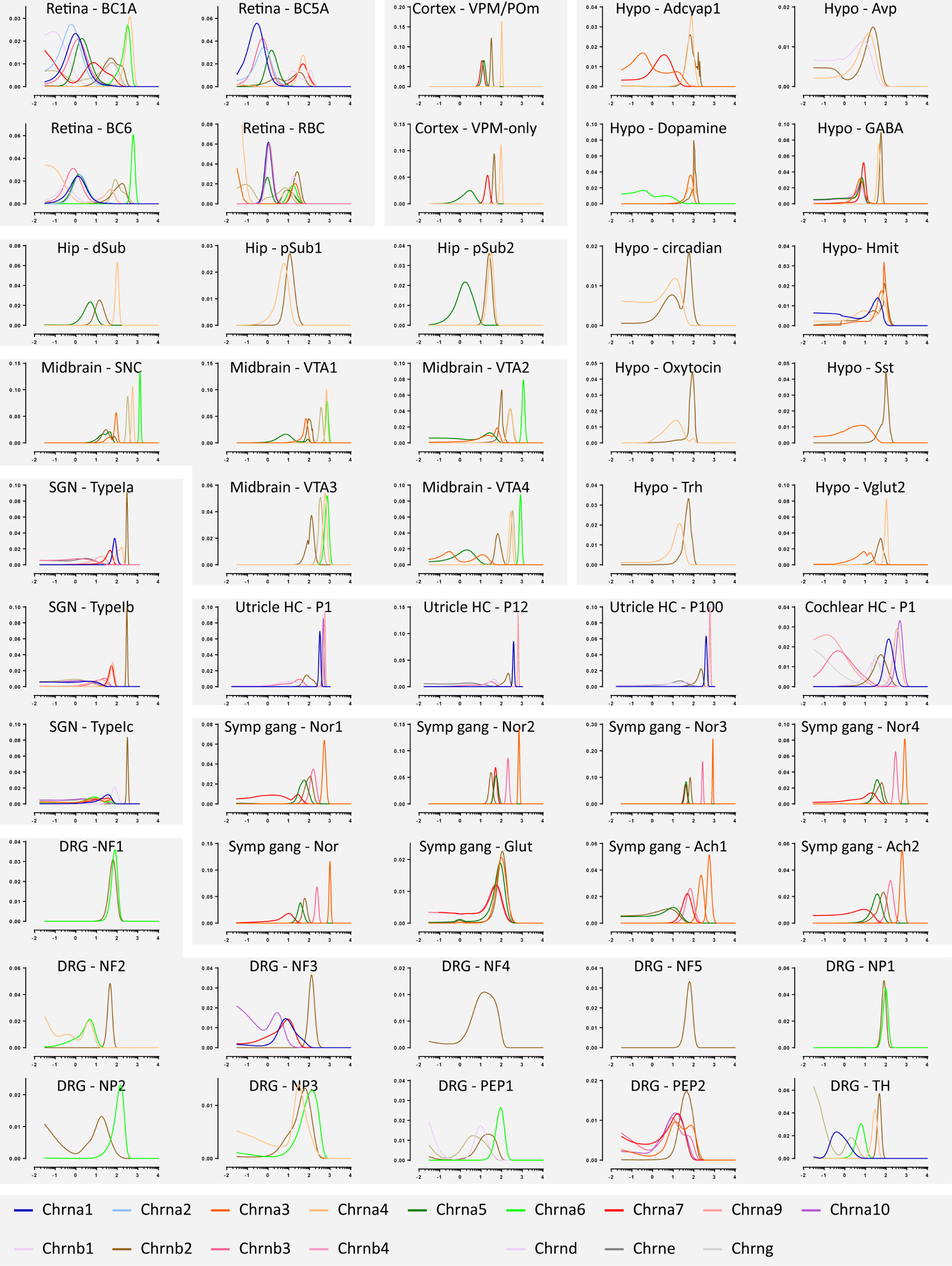
Joint posterior probabilities of gene expression levels inferred for the nAChR subunits expressed in the cell types and datasets analysed (see SI Methods for details).

**Supplementary Figure 6.**
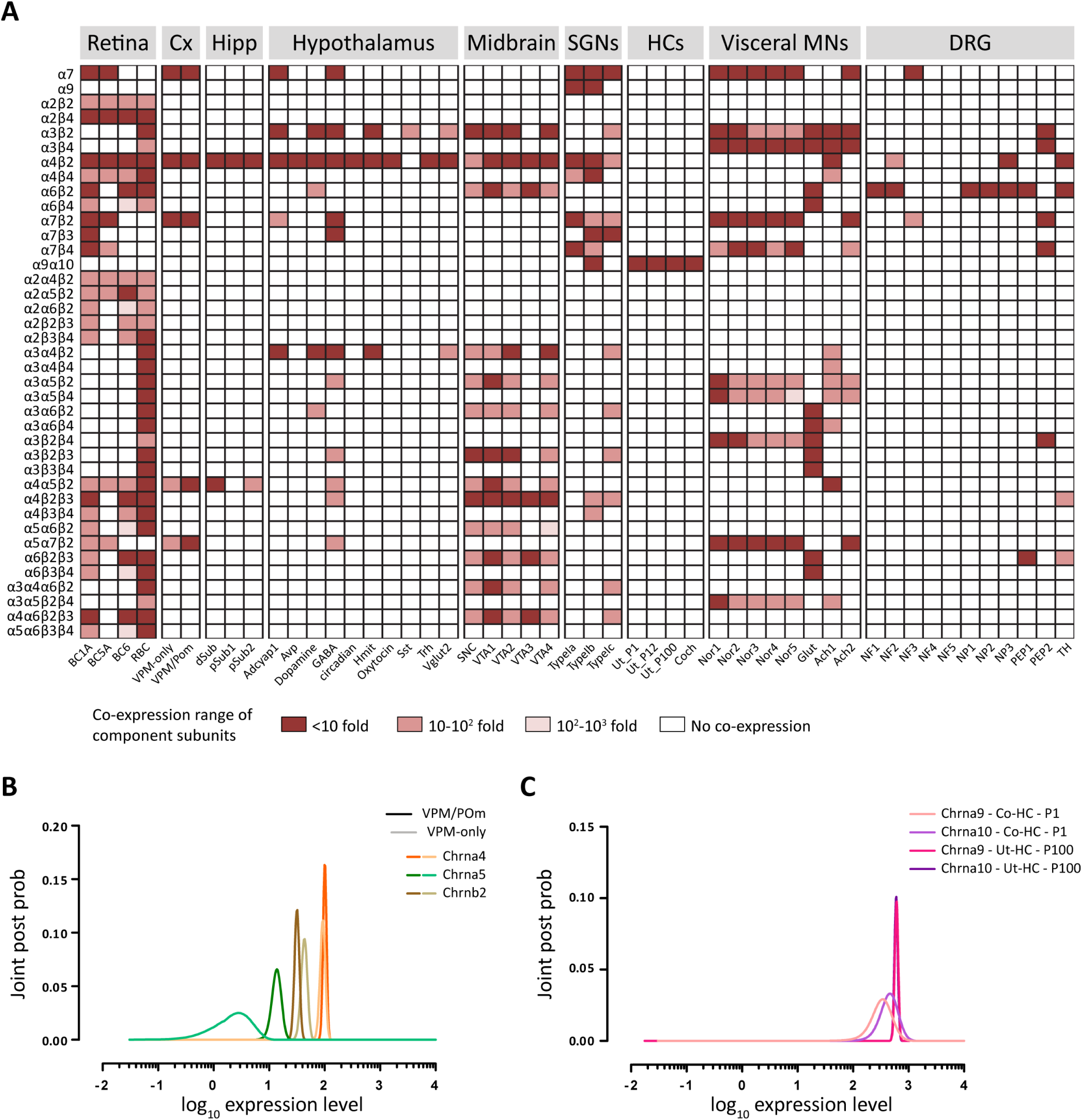
**A.** Co-expression of subunits comprising known nAChR assemblies. Dark red squares, all component subunits are co-expressed within a 10-fold range of expression level. Light red squares, all component subunits are co-expressed within a 100-fold range of expression level. Pink squares, all component subunits are expressed within a 1000-fold range of expression level. White squares, at least one subunit of that receptor assembly in not expressed in that cell type. **B.** Estimated joint posterior probability distributions for expression levels of Chrna4, ChrnaS and Chrnb2 in VPM-only and VPM/POm projecting layer VI neurons. **C.** Estimated joint posterior probability distributions for expression levels of Chrna9 and Chrnal0 in inner ear cochlear hair cells (Pl) and utricle hair cells (Pl00).

**Supplementary Figure 7.**
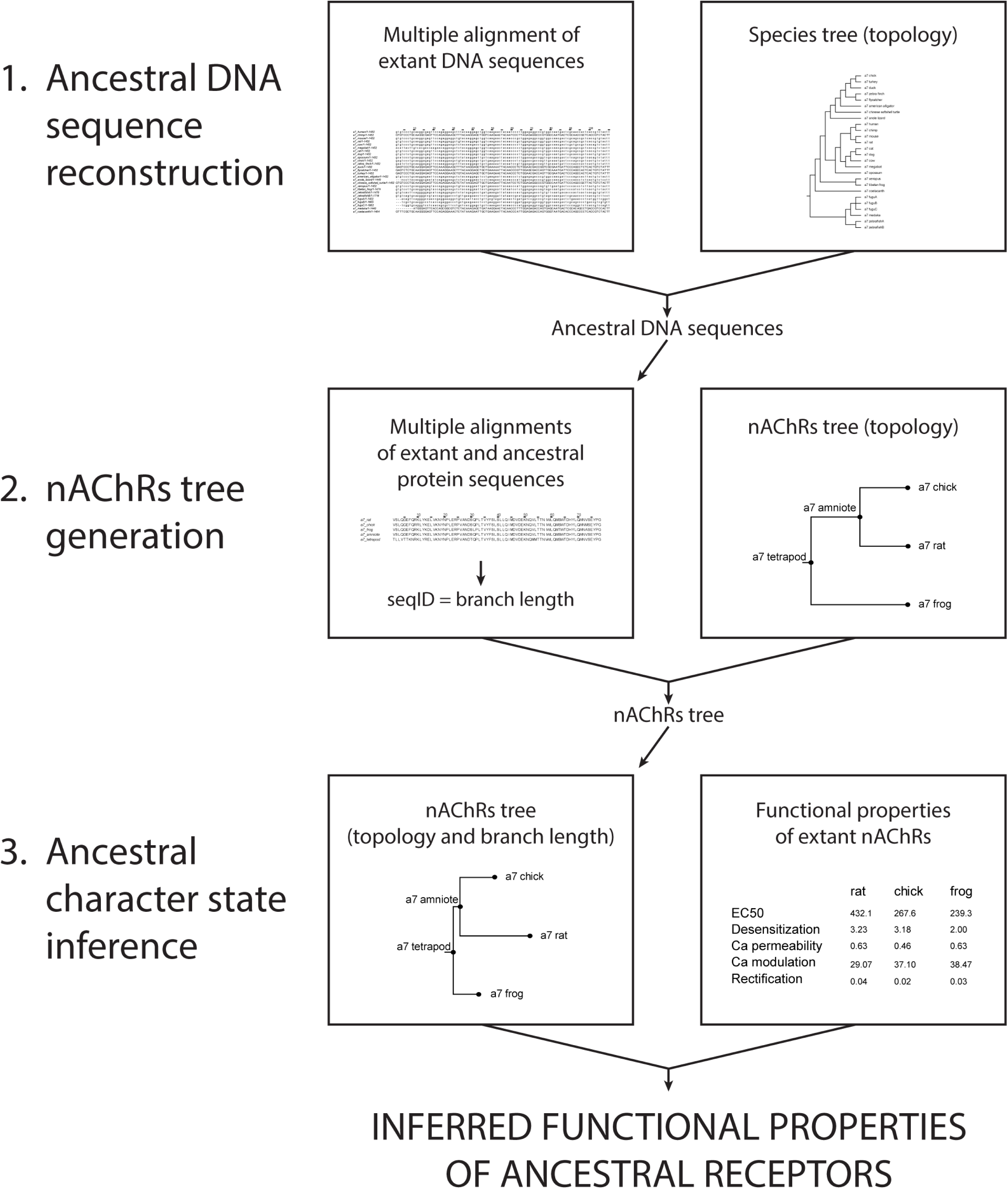
Experimental workflow detailing the steps followed for the inference of ancestral character states of the functional properties of tetrapod and amniote ancestral α4β2, α7 and α9α10 receptors.

**Supplementary Figure 8.**
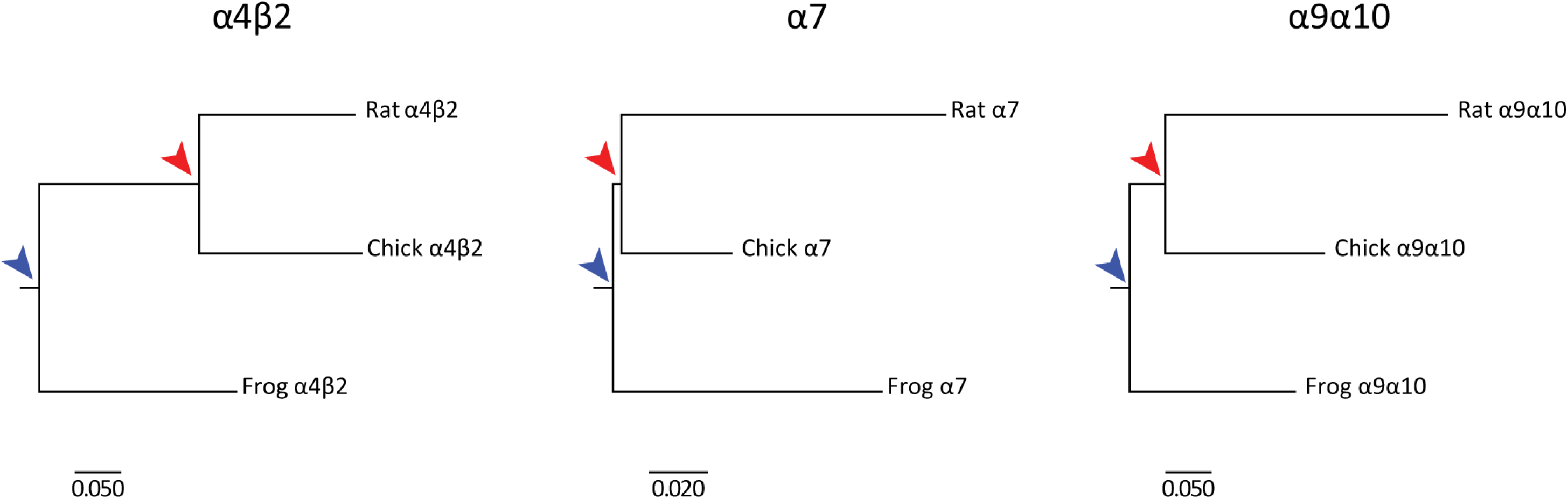
nAChRs trees used for the inference of ancestral state states. The tree topology corresponds to the species phylogenetic relationship. The branch lengths correspond to those determined from aminoacid sequence identity analysis between rat-amniote, chick-amniote, amniote-tetrapod and frog-tetrapod pairs of subunits. For the heteromeric α4β2 receptors, branch lengths were calculated assuming a (α4)2(β2)3 assembly. For the heteromeric α9α10 receptors branch lengths were calculated assuming a (α9)2(α10)3 stoichiometry (Plazas et al., 2005). Blue arrows, tetrapod ancestor. Red arrows, amniote ancestor.

**Supplementary Figure 9.**
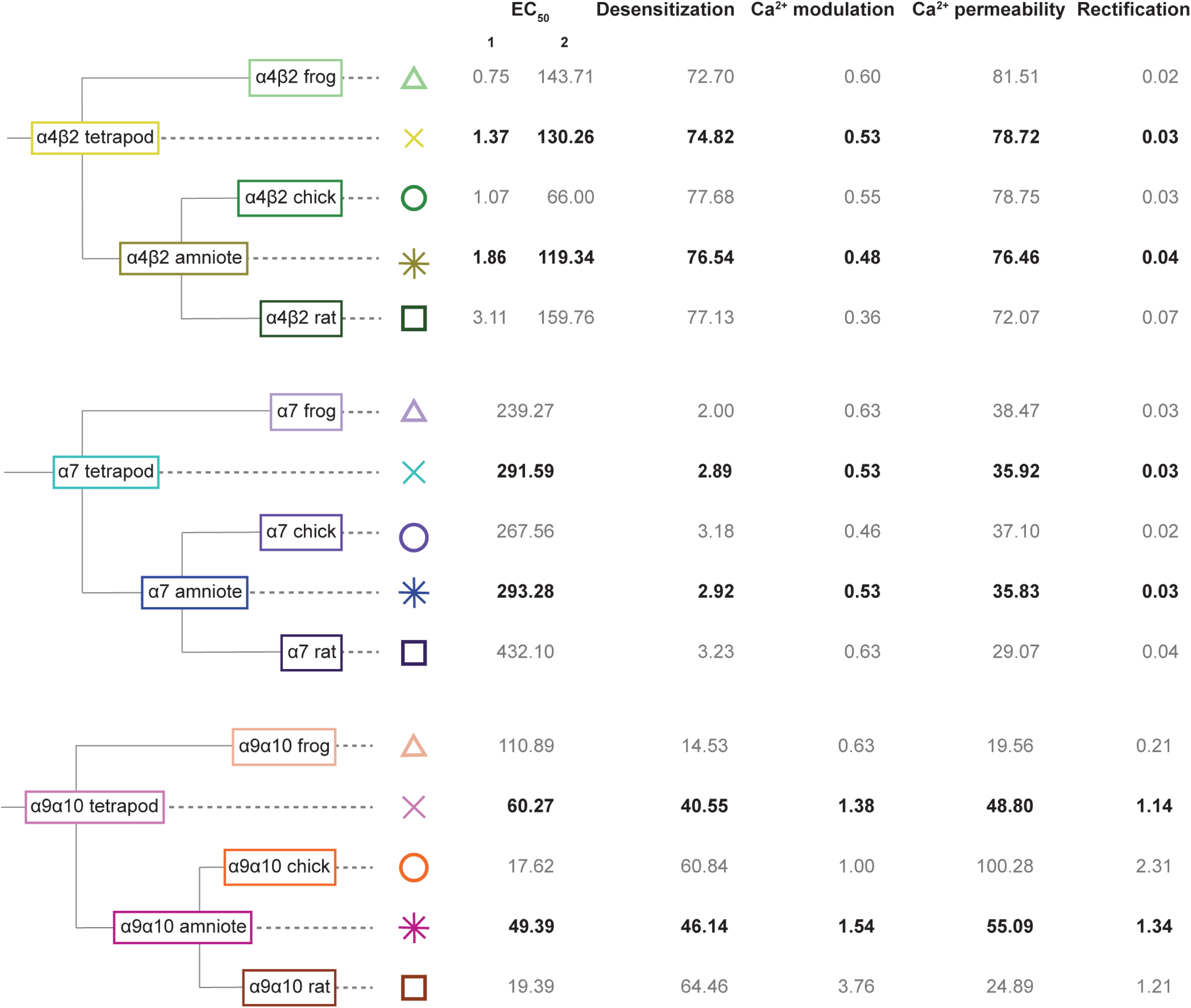
Inferred ancestral character states for amniote and tetrapod α9α10, α4β2 and α7 nAChRs. The biophysical properties of ancestral receptors inferred by maximum likelihood are shown in black. The values in grey correspond to the biophysical properties of extant receptors determined experimentally (Table S7) and used as input for the inference of ancestral character state. The symbols next to each receptor correspond to the symbols in Figure 8.

**Table S1.**
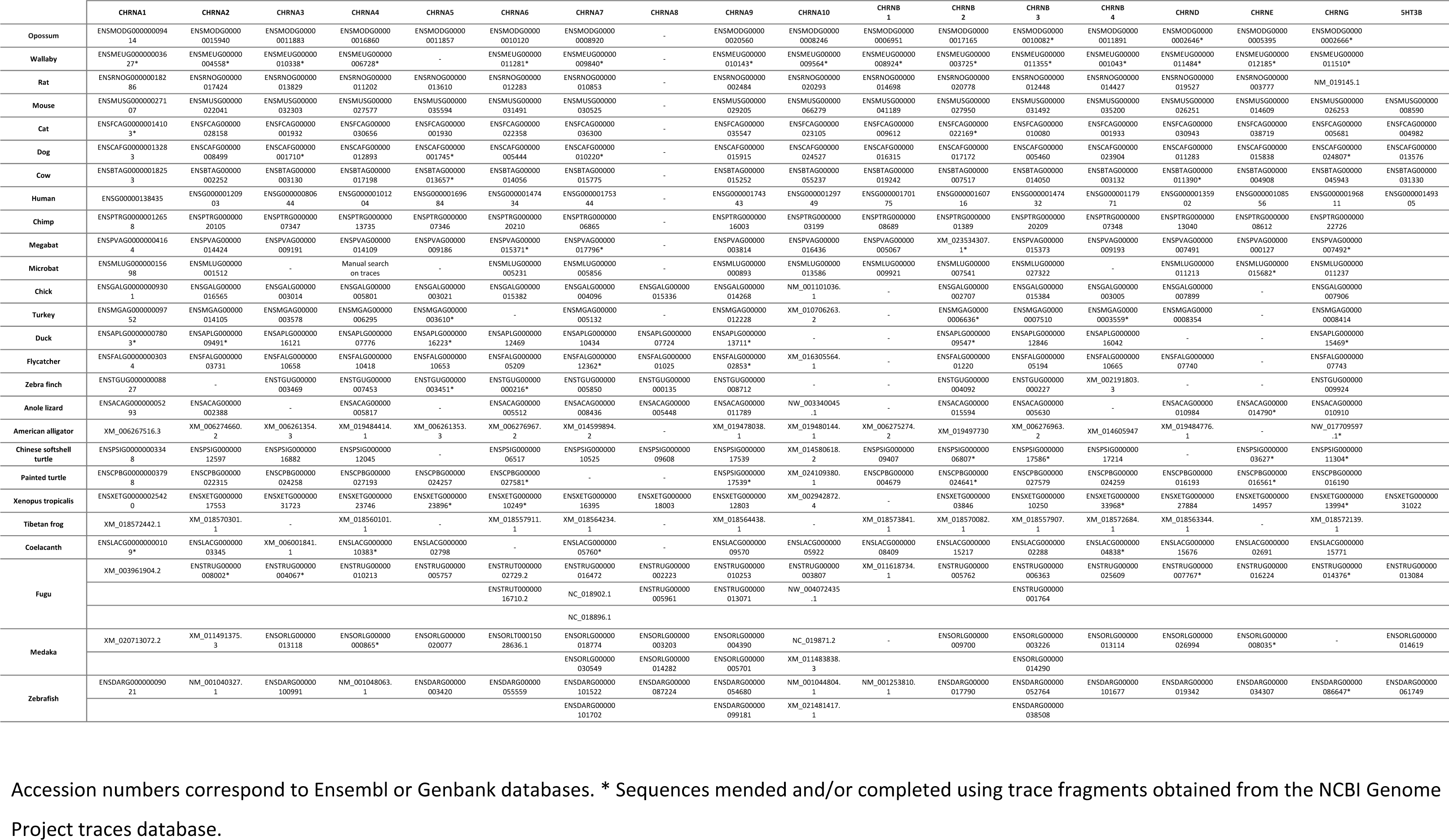
Accession numbers for sequences used in the phylogenetic analysis.

**Table S2.**
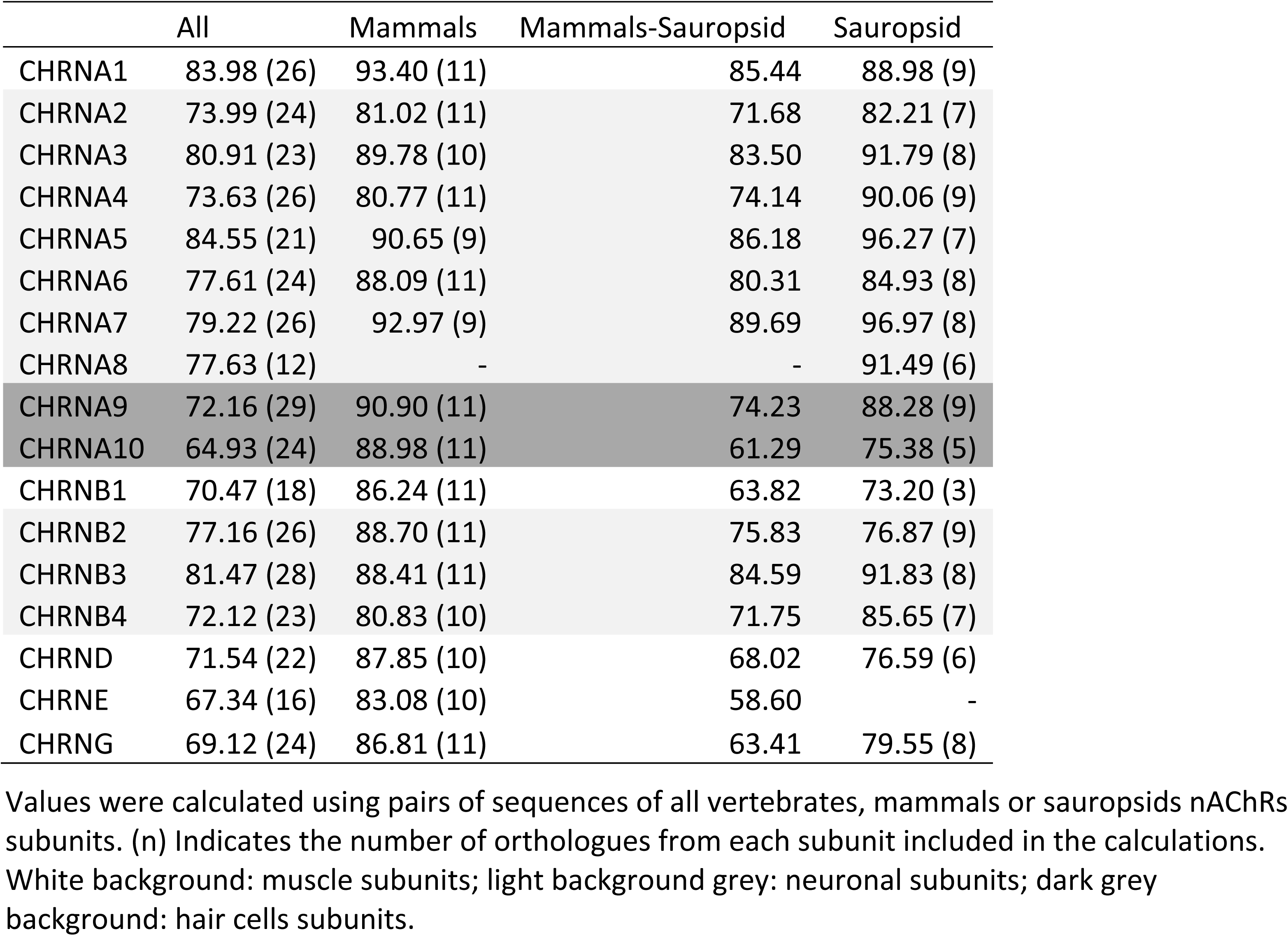
Mean percentage sequence identity (%seqID).

**Table S3.**
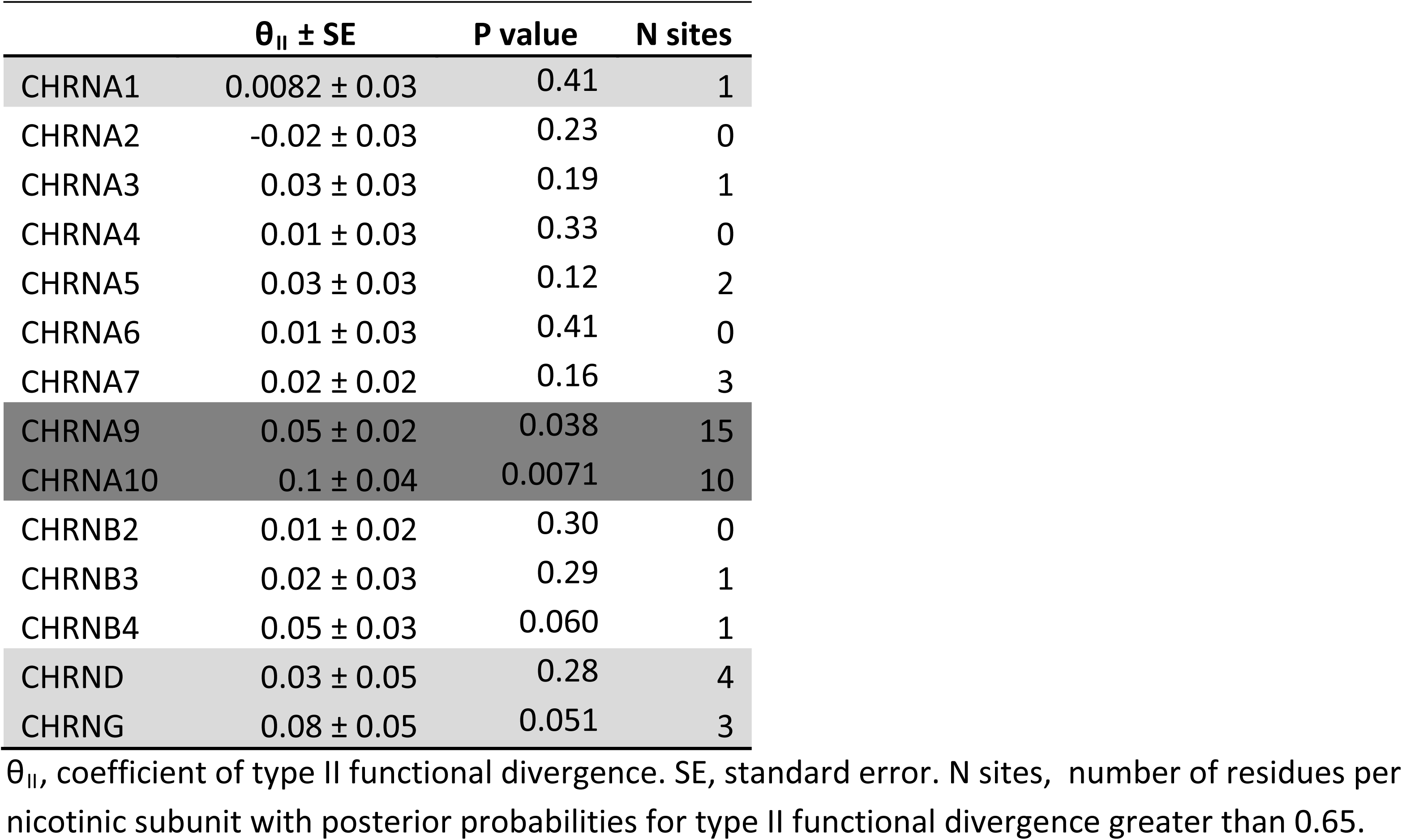
Type II functional divergence coefficients for nAChR subunits

**Table S4.**
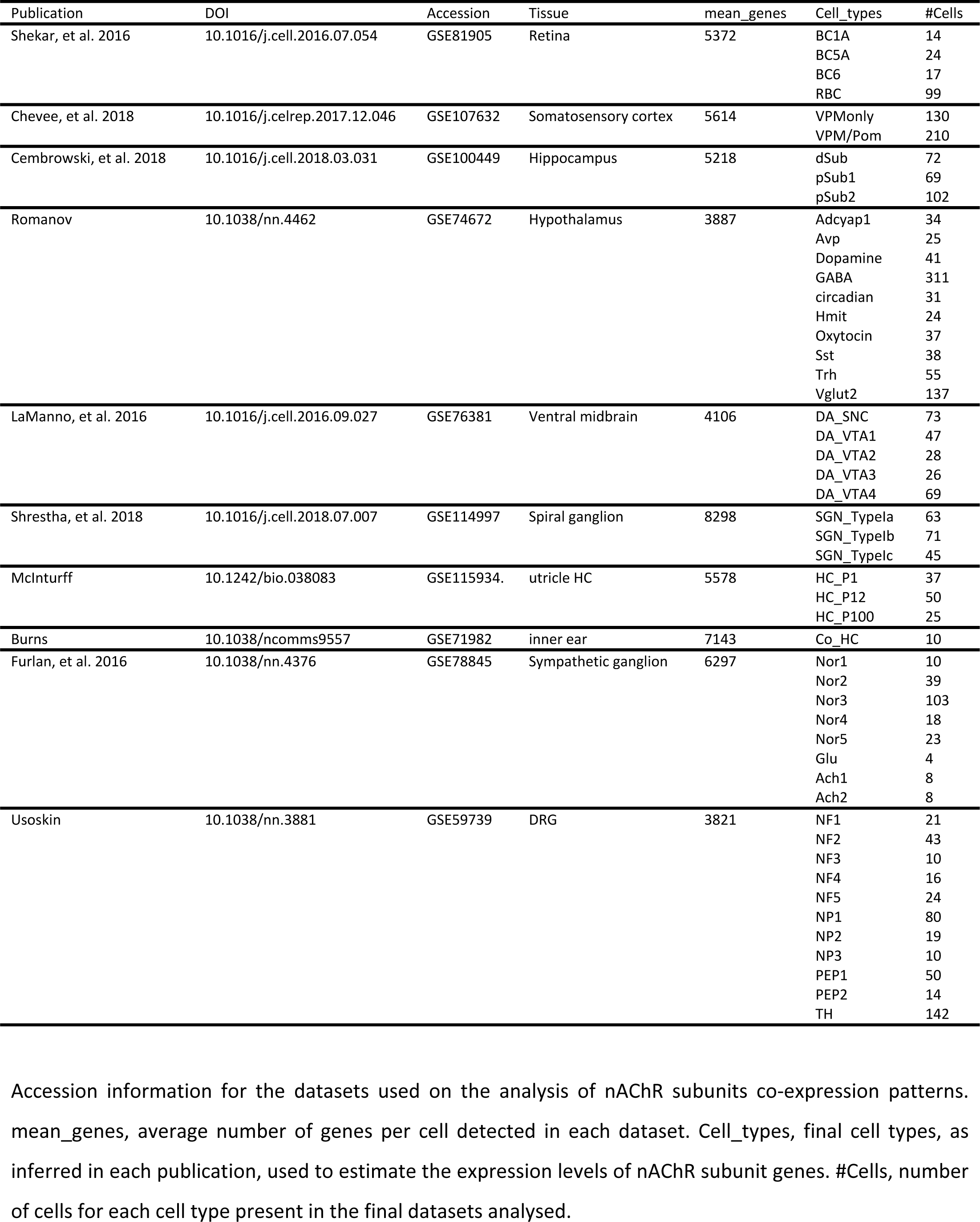
Single-cell RNA sequencing datasets.

**Table S5.**
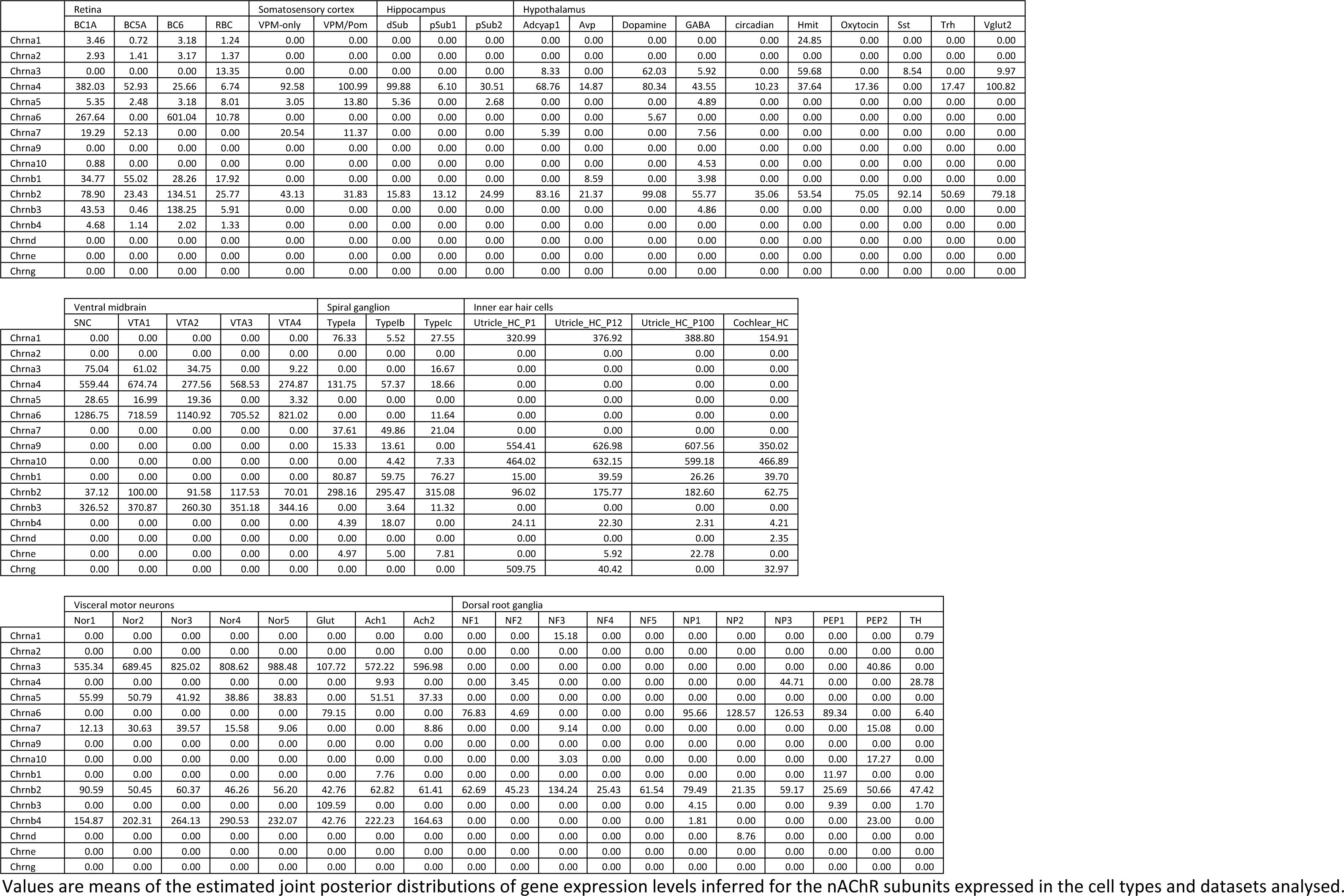
Mean expression of nAChR subunits genes across cell types.

**Table S6.** Neuronal nAChRs experimentally validated assemblies.

| nAChR subtype | DOI | Reference |
| --- | --- | --- |
| $\alpha 7$ | 10.1016/0896-6273(90)90344-F | Couturier et al., 1990 |
| $\alpha 8$ | PMID: 7509438 | Gerzanich et al., 1994 |
| $\alpha 9$ | 10.1016/0092-8674(94)90555-X | Elgoyhen 1994 et al., 1994 |
| $\alpha 10$ | 10.1093/molbev/msu258 | Lipovsek et al., 2014 |
| $\alpha 2\beta 2$ | 10.1016/0896-6273(88)90208-5 | Deneris et al., 1988 |
| $\alpha 2\beta 4$ | 10.1016/0896-6273(89)90207-9 | Duvoisin et al., 1989 |
| $\alpha 3\beta 2$ | 10.1073/pnas.84.21.7763 | Boulter et al., 1987 |
| $\alpha 3\beta 4$ | 10.1016/0896-6273(89)90207-9 | Duvoisin et al., 1989 |
| $\alpha 4\beta 2$ | 10.1073/pnas.84.21.7763 | Boulter et al., 1987 |
| $\alpha 4\beta 4$ | 10.1016/0896-6273(89)90207-9 | Duvoisin et al., 1989 |
| $\alpha 6\beta 2$ | 10.1046/j.1460-9568.1998.00001.x | Fucile et al., 1998 |
| $\alpha 6\beta 4$ | 10.1124/mol.106.027326 | Tumkosit et al., 2006 |
| $\alpha 7\beta 2$ | 10.1113/jphysiol.2001.013847 | Khiroug et al., 2002 |
| $\alpha 7\beta 3$ | 10.1074/jbc.274.26.18335 | Palma et al., 1999 |
| $\alpha 7\beta 4$ | 110.1111/j.1471-4159.2012.07931.x | Criado et al., 2012 |
| $\alpha 9\alpha 10$ | 10.1073/pnas.051622798 | Elgoyhen et al., 2001 |
| $\alpha 2\alpha 5\beta 2$ | 10.1124/mol.63.6.1329 | Vailati et al., 2003 |
| $\alpha 2\alpha 6\beta 2$ | 10.1124/mol.105.015925 | Gotti et al., 2005 |
| $\alpha 2\beta 2\beta 3$ | 10.1111/j.1471-4159.2012.07685.x | Dash et al., 2012 |
| $\alpha 2\beta 3\beta 4$ | 10.1111/j.1471-4159.2012.07685.x | Dash et al., 2012 |
| $\alpha 3\alpha 4\beta 2$ | 10.1523/JNEUROSCI.2112-05.2005 | Turner et al., 2005 |
| $\alpha 3\alpha 4\beta 4$ | 10.1523/JNEUROSCI.2112-05.2005 | Turner et al., 2005 |
| $\alpha 3\alpha 5\beta 2$ | 10.1074/jbc.271.30.17656 | Wang et al., 1996 |
| $\alpha 3\alpha 5\beta 4$ | 10.1074/jbc.271.30.17656 | Wang et al., 1996 |
| $\alpha 3\alpha 6\beta 2$ | 10.1016/S0028-3908(00)00144-1 | Kuryatov et al., 2000 |
| $\alpha 3\alpha 6\beta 4$ | 10.1046/j.1460-9568.1998.00001.x | Fucile et al., 1998 |
| $\alpha 3\beta 2\beta 4$ | 10.1523/JNEUROSCI.2112-05.2005 | Turner et al., 2005 |
| $\alpha 3\beta 2\beta 3$ | 10.1111/j.1471-4159.2012.07685.x | Dash et al., 2012 |
| $\alpha 3\beta 3\beta 4$ | 10.1074/jbc.273.25.15317 | Groot-Kormelink et al., 1998 |
| $\alpha 4\alpha 5\beta 2$ | 10.1038/380347a0 | Ramirez-Latorre et al., 1996 |
| $\alpha 4\beta 2\beta 3$ | 10.1124/mol.108.046789 | Kuryatov et al., 2008 |
| $\alpha 4\beta 3\beta 4$ | 10.1111/j.1471-4159.2012.07685.x | Dash et al., 2012 |
| $\alpha 5\alpha 6\beta 2$ | 10.1016/S0028-3908(00)00144-1 | Kuryatov et al., 2000 |
| $\alpha 5\alpha 7\beta 2$ | 10.1111/j.1749-6632.1999.tb11331.x | Girod et al., 1999 |
| $\alpha 6\beta 2\beta 3$ | 10.1124/mol.106.027326 | Tumkosit et al., 2006 |
| $\alpha 6\beta 3\beta 4$ | 10.1016/S0028-3908(00)00144-1 | Kuryatov et al., 2000 |
| $\alpha 3\alpha 4\alpha 6\beta 2$ | 10.1111/j.1471-4159.2008.05282.x | Cox et al., 2008 |
| $\alpha 3\alpha 5\beta 2\beta 4$ | 10.1074/jbc.270.9.4424 | Conroy and Berg, 1995 |
| $\alpha 4\alpha 6\beta 2\beta 3$ | 10.1124/mol.110.066159 | Kuryatov and Lindstrom, 2010 |
| $\alpha 5\alpha 6\beta 3\beta 4$ | 10.1124/jpet.104.075069 | Grinevich et al., 2005 |
| $\alpha 1\beta 1\gamma \delta$ | 10.1038/321406a0 | Mishina et al., 1986 |
| $\alpha 1\beta 1\delta \epsilon$ | 10.1038/321406a0 | Mishina et al., 1986 |

Experimentally validated pentameric combinations of nAChR subunits and relevant literature references.

**Table S7.**
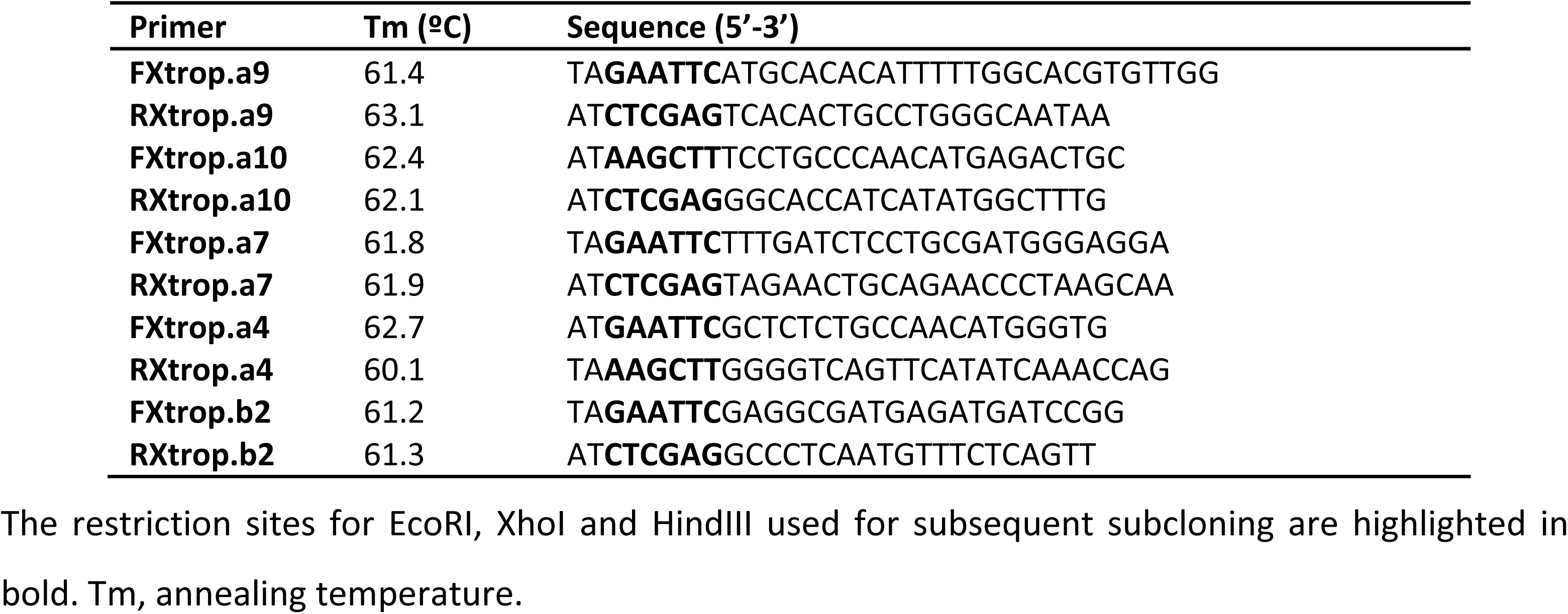
Primers used for the cloning of *X. tropicalis* nAChRs subunits.

**Table S8.** Biophysical properties and statistical comparisons from rat, chicken and frog α4β2, α7 and α9α10 receptors.

| Receptor |  | α4β2 |  |  |  |  |  |
| --- | --- | --- | --- | --- | --- | --- | --- |
| Species |  | Rat |  | Chicken |  | Frog |  |
| ACh sensitivity | Mean ± S.E.M (n) | 3.11 ± 2.11 (3) | 159.76 ± 35.67 (4) | 1.07 ± 0.15 (6) | 66.00 ± 9.76 (4) | 0.75 ± 0.02 (7) | 143.71 ± 51.89 (5) |
|  | ANOVA (p-value) | EC50_1: Rat vs Chick = 0.1346, Rat vs Frog = 0.1018, Chick vs Frog = 0.6927<sup>P</sup> |  |  |  |  |  |
|  |  | EC50_2: Rat vs Chick = 0.3743, Rat vs Frog = 0.7809, Chick vs Frog = 0.3743<sup>P</sup> |  |  |  |  |  |
| Desensitization | Mean ± S.E.M (n) | 77.13 ± 1.21 (4) |  | 77.68 ± 5.16 (7) |  | 72.70 ± 3.04 (10) |  |
|  | ANOVA (p-value) | Rat vs Chick = 0.9336, Rat vs Frog = 0.7326, Chick vs Frog = 0.7204<sup>P</sup> |  |  |  |  |  |
| Ca<sup>2+</sup> Modulation | Mean ± S.E.M (n) | 0.36 ± 0.04 (5) |  | 0.55 ± 0.03 (9) |  | 0.60 ± 0.08 (5) |  |
|  | ANOVA (p-value) | Rat vs Chick > 0.999, Rat vs Frog > 0.9999, Chick vs Frog > 0.9999<sup>P</sup> |  |  |  |  |  |
| Ca<sup>2+</sup> Permeability | Mean ± S.E.M (n) | 72.07 ± 9.62 (4) |  | 78.75 ± 5.04 (6) |  | 81.51 ± 7.91 (6) |  |
|  | ANOVA (p-value) | Rat vs Chick = 0.7999, Rat vs Frog = 0.7892, Chick vs Frog = 0.7999<sup>P</sup> |  |  |  |  |  |
| Rectification profile | Mean ± S.E.M (n) | 0.07 ± 0.03 (5) |  | 0.03 ± 0.03 (6) |  | 0.02 ± 0.03 (6) |  |
|  | ANOVA (p-value) | Rat vs Chick = 0.6838, Rat vs Frog = 0.4995, Chick vs Frog = 0.9437<sup>P</sup> |  |  |  |  |  |
| Receptor |  | α7 |  |  |  |  |  |
| Species |  | Rat |  | Chicken |  | Frog |  |
| ACh sensitivity | Mean ± S.E.M (n) | 432.1 ± 123.45 (8) |  | 267.56 ± 87.98 (8) |  | 239.27 ± 33.90 (7) |  |
|  | ANOVA (p-value) | Rat vs Chick = 0.365, Rat vs Frog = 0.6723, Chick vs Frog > 0.9999<sup>NP</sup> |  |  |  |  |  |
| Desensitization | Mean ± S.E.M (n) | 3.23 ± 0.79 (6) |  | 3.18 ± 0.54 (9) |  | 2.00 ± 0.41 (6) |  |
|  | ANOVA (p-value) | Rat vs Chick > 0.9999, Rat vs Frog = 0.3743, Chick vs Frog = 0.2496<sup>NP</sup> |  |  |  |  |  |
| Ca<sup>2+</sup> Modulation | Mean ± S.E.M (n) | 0.63 ± 0.10 (4) |  | 0.46 ± 0.06 (4) |  | 0.63 ± 0.08 (4) |  |
|  | ANOVA (p-value) | Rat vs Chick > 0.999, Rat vs Frog > 0.9999, Chick vs Frog > 0.9999<sup>P</sup> |  |  |  |  |  |
| Ca<sup>2+</sup> Permeability | Mean ± S.E.M (n) | 29.07 ± 7.68 (4) |  | 37.10 ± 11.82 (4) |  | 38.47 ± 3.98 (7) |  |
|  | ANOVA (p-value) | Rat vs Chick = 0.7447, Rat vs Frog = 0.7447, Chick vs Frog = 0.8931<sup>P</sup> |  |  |  |  |  |
| Rectification profile | Mean ± S.E.M (n) | 0.04 ± 0.02 (5) |  | 0.02 ± 0.003 (5) |  | 0.03 ± 0.01 (5) |  |
|  | ANOVA (p-value) | Rat vs Chick = 0.5909, Rat vs Frog = 0.9706, Chick vs Frog = 0.7295<sup>P</sup> |  |  |  |  |  |
| Receptor |  | α9α10 |  |  |  |  |  |
| Species |  | Rat |  | Chicken |  | Frog |  |
| ACh sensitivity | Mean ± S.E.M (n) | 19.39 ± 2.08 (9) |  | 17.62 ± 3.33 (6) |  | 110.89 ± 25.00 (7) |  |
|  | ANOVA (p-value) | Rat vs Chick > 0.9999, Rat vs Frog = 0.0026<sup>**</sup>, Chick vs Frog = 0.006<sup>**NP</sup> |  |  |  |  |  |
| Desensitization | Mean ± S.E.M (n) | 64.46 ± 3.65 (5) |  | 60.84 ± 4.23 (9) |  | 14.53 ± 3.01 (7) |  |
|  | ANOVA (p-value) | Rat vs Chick > 0.9999, Rat vs Frog = 0.0051<sup>**</sup>, Chick vs Frog = 0.0042<sup>**NP</sup> |  |  |  |  |  |
| Ca<sup>2+</sup> Modulation | Mean ± S.E.M (n) | 3.76 ± 0.73 (5) |  | 1.00 ± 0.07 (4) |  | 0.63 ± 0.05 (12) |  |
|  | ANOVA (p-value) | Rat vs Chick < 0.0001<sup>****</sup>, Rat vs Frog < 0.0001<sup>****</sup>, Chick vs Frog > 0.9981<sup>P</sup> |  |  |  |  |  |
| Ca<sup>2+</sup> Permeability | Mean ± S.E.M (n) | 24.89 ± 2.81 (6) |  | 100.28 ± 14.02 (6) |  | 19.56 ± 3.38 (10) |  |
|  | ANOVA (p-value) | Rat vs Chick = 0.0299<sup>*</sup>, Rat vs Frog > 0.9999, Chick vs Frog = 0.0013<sup>**NP</sup> |  |  |  |  |  |
| Rectification profile | Mean ± S.E.M (n) | 1.21 ± 0.07 (11) |  | 2.31 ± 0.34 (10) |  | 0.21 ± 0.05 (8) |  |
|  | ANOVA (p-value) | Rat vs Chick = 0.0229<sup>*</sup>, Rat vs Frog = 0.0406<sup>*</sup>, Chick vs Frog < 0.0001<sup>****NP</sup> |  |  |  |  |  |

Values are shown as mean ± S.E.M. (n). <sup>P</sup>: parametric, <sup>NP</sup>: non-parametric analysis. ACh sensitivity: EC<sub>50</sub> value; Desensitization rate: percentage current remaining 20 seconds (5 sec for $\alpha 7$ receptors) after peak response to a 10-fold concentration of EC<sub>50</sub> ACh; Ca<sup>2+</sup> Modulation: current elicited by EC<sub>50</sub> ACh at Ca<sup>2+</sup> 0.5 mM relative to Ca<sup>2+</sup> 3 mM; Ca<sup>2+</sup> Permeability: percentage of remaining current after BAPTA-AM treatment; Rectification profile: current recorded at +40 mV relative to that recorded at -90 mV.

**Table S9.** Normalized biophysical properties

| Receptor | ACh sensitivity<br>(EC<sub>50</sub>) | Desensitization<br>(%/ after ACh peak) | Ca<sup>2+</sup> modulation<br>(I<sub>0.5 mM</sub>/I<sub>3 mM</sub>) | Ca<sup>2+</sup> permeability<br>(%/ post-BAPTA) | Rectification<br>(I<sub>+40 mV</sub>/I<sub>-90 mV</sub>) |
| --- | --- | --- | --- | --- | --- |
| Rat $\alpha 9\alpha 10$ | 0.045 | 0.830 | 1.000 | 0.248 | 0.525 |
| Chick $\alpha 9\alpha 10$ | 0.041 | 0.783 | 0.266 | 1.000 | 1.000 |
| Frog $\alpha 9\alpha 10$ | 0.257 | 0.187 | 0.167 | 0.195 | 0.091 |
| Rat $\alpha 4\beta 2$ | 0.007 | 0.993 | 0.097 | 0.719 | 0.029 |
| Chick $\alpha 4\beta 2$ | 0.002 | 1.000 | 0.146 | 0.785 | 0.014 |
| Frog $\alpha 4\beta 2$ | 0.002 | 0.936 | 0.158 | 0.813 | 0.008 |
| Rat $\alpha 7$ | 1.000 | 0.042 | 0.167 | 0.290 | 0.016 |
| Chick $\alpha 7$ | 0.619 | 0.041 | 0.121 | 0.370 | 0.009 |
| Frog $\alpha 7$ | 0.554 | 0.026 | 0.168 | 0.384 | 0.015 |
| Amniote $\alpha 4\beta 2$ | 0.004 | 0.985 | 0.128 | 0.762 | 0.019 |
| Tetrapod $\alpha 4\beta 2$ | 0.003 | 0.963 | 0.141 | 0.785 | 0.014 |
| Amniote $\alpha 7$ | 0.679 | 0.038 | 0.141 | 0.357 | 0.012 |
| Tetrapod $\alpha 7$ | 0.675 | 0.037 | 0.142 | 0.358 | 0.012 |
| Amniote $\alpha 9\alpha 10$ | 0.114 | 0.594 | 0.409 | 0.549 | 0.581 |
| Tetrapod $\alpha 9\alpha 10$ | 0.139 | 0.522 | 0.366 | 0.487 | 0.494 |

Normalized biophysical properties from extant receptors used in PCA analysis (top) and normalised inferred biophysical properties from ancestral receptors. Values are normalised relative to the maximum obtained for each parameter.

**Table S10.** PCA analysis of functional properties.

|  | PC1 | PC2 | PC3 | PC4 | PC5 |
| --- | --- | --- | --- | --- | --- |
| **ACh sensitivity ($EC_{50}$)** | 0.55 | 0.04 | 0.35 | 0.72 | 0.23 |
| **Desensitization (%I after ACh peak)** | -0.56 | -0.11 | -0.35 | 0.41 | 0.62 |
| **Ca<sup>2+</sup> modulation ($I_{0.5\text{ mM}}/I_{3\text{ mM}}$)** | -0.18 | 0.77 | -0.26 | 0.36 | -0.42 |
| **Ca<sup>2+</sup> permeability (%I post-BAPTA)** | -0.46 | -0.46 | 0.37 | 0.38 | -0.55 |
| **Rectification ($I_{+40\text{ mV}}/I_{-90\text{ mV}}$)** | -0.36 | 0.42 | 0.74 | -0.21 | 0.31 |
| **Proportion of the variability** | 0.54 | 0.28 | 0.14 | 0.03 | 0.006 |
| **Accumulated proportion** | 0.54 | 0.82 | 0.96 | 0.99 | 1.00 |

*Upper panel:* loading of each experimental parameter on the five principal components. *Lower panel:* proportion and accumulated proportion of the variability represented by each principal component.

**Table S11.** Estimated distances between extant and inferred ancestral α9α10, α4β2 and α7 nAChRs based on protein sequence identity.

| **$\alpha 4\beta 2$** |  |  |
| --- | --- | --- |
| **Comparison** | **seqID** | **1-seqID** |
| Rat-amniote | 0.824 | 0.176 |
| Chick-amniote | 0.816 | 0.184 |
| Amniote-tetrapod | 0.819 | 0.181 |
| Frog-tetrapod | 0.777 | 0.223 |
| **$\alpha 7$** |  |  |
| **Comparison** | **seqID** | **1-seqID** |
| Rat-amniote | 0.888 | 0.112 |
| Chick-amniote | 0.962 | 0.038 |
| Amniote-tetrapod | 0.997 | 0.003 |
| Frog-tetrapod | 0.907 | 0.093 |
| **$\alpha 9\alpha 10$** |  |  |
| **Comparison** | **seqID** | **1-seqID** |
| Rat-amniote | 0.683 | 0.317 |
| Chick-amniote | 0.821 | 0.179 |
| Amniote-tetrapod | 0.96 | 0.04 |
| Frog-tetrapod | 0.814 | 0.186 |

## Supplementary File Legends

**Supplementary File 1. Multiple alignment of nAChRs subunits.** The alignment includes 392 nAChR subunits sequences from 29 different species together with 9 sequences from 5HT3B vertebrate subunits (outgroup).

**Supplementary File 2. Aminoacid sequence identities.** Values for all pairwise comparisons for all nAChR subunits from amniotes.

**Supplementary File 3. Type II functional divergence analysis using DIVERGE 3.0.** For each nAChR subunit analysed, the main tree, cluster-mammalian tree and cluster-sauropsid tree are shown in parenthetical notation. Theta-II values and standard error (SE) were calculated by comparing the designated clusters. Posterior probabilities (Pp) per site were calculated from the Posterior ratio (Pr) values obtained from the type II function by *Pp = Pr / (1+Pr)*, as in Gu et al, 2006.

**Supplementary File 4. Sequences used for the reconstruction of ancestral nAChR subunits.** DNA sequences from α4, α7, α9, α10 and β2 orthologues used to reconstruct DNA sequences from amniote and tetrapod ancestors. The species trees, in parenthetic format, are shown before the sequence alignment, in fasta format. The branch lengths in the species trees were inferred from each alignment and indicate the amount of accumulated changes.

**Supplementary File 5. Extant and inferred ancestral nAChRs protein sequences.** Protein sequences from rat, chicken, frog, amniote ancestor and tetrapod ancestor, from α7, (α4)_2_(β2)_3_ and (α9)_2_(α10)_3_ nAChRs were aligned and the sequence identity between pairs of receptors was used to calculate the branch lengths assigned to the receptor trees (Supplementary Figure 6) used to infer ancestral character states of biophysical properties (Supplementary Figure 7).

